# A long non-coding RNA is a key factor in the evolution of insect eusociality

**DOI:** 10.1101/2022.05.10.491402

**Authors:** Carlos A. M. Cardoso-Junior, Gustavo J. Tibério, Denyse C. Lago, Luiz Carlos Vieira, José C. Rosa, Alexandre R. Paschoal, Isobel Ronai, Benjamin P. Oldroyd, Klaus Hartfelder

**Affiliations:** Departamento de Biologia Celular e Molecular, Faculdade de Medicina de Ribeirão Preto, Universidade de São Paulo, Ribeirão Preto, SP, Brazil.; Departamento de Genética, Faculdade de Medicina de Ribeirão Preto, Universidade de São Paulo, Ribeirão Preto, SP, Brazil.; Universidade Tecnológica Federal do Paraná, Cornélio Procópio, PR, Brazil.; Behaviour, Ecology and Evolution (BEE) laboratory, Macleay Building A12, University of Sydney, Sydney NSW 2006, Australia.; Wissenschaftskolleg zu Berlin, Wallotstrasse 19 D-14193, Berlin, Germany.

**Author notes:** These authors contributed equally (Carlos A. M. Cardoso-Junior, Gustavo J. Tibério). **AUTHOR CONTRIBUTIONS** CAM designed, conceived, performed and analyzed the experiments on adult bees, helped in the pulldown assay, cloned and sequenced the full-length *lncov1* sequence, designed the RNA-seq analyses together with LCV and wrote the manuscript draft together with KH. GJT conceived, designed the study, performed and analyzed the larval experiments. DCL performed the JH treatment experiment, analyzed FISH assays together with GJT and quantified the caste-specific *Tudor-SN* gene expression. LCV performed the bioinformatic analyses for RNA-seq datasets. JCR and GJT performed the mass spectrometry analysis. IR helped in gene expression analyses of adult honey bees and in the setting up of the cages for the diet and QMP treatments. BPO designed the experiments with adult bees together with CAM, performed social manipulations in field colonies, and supervised the study. ARP performed the bioinformatics analysis on the conservation of the *lncov1* gene. KH conceived, designed, supervised the study, and, together with CAM drafted the manuscript. All authors contributed critically to this paper and approved the final version.

**Keywords:** Honeybees, reproductive division of labor, queen mandibular pheromone (QMP), lncRNA, royal jelly, epigenetics

## Abstract

Insect sociality is a major evolutionary transition based on the suppression of worker reproduction in favor of the reproductive monopoly of the queen. In the honey bee (*Apis mellifera*) model organism, the development of the two female caste phenotypes, queen and worker, is triggered by differences in their larval diets. However, the mechanistic details underlying their respective developmental trajectories, as well as the maintenance of sterility in the adult workers, are still not fully understood. Here we show that the long non-coding RNA *lncov1* interacts with the Tudor staphylococcus nuclease (Tudor-SN) protein to form a regulatory module that promotes apoptosis in the ovaries of worker larvae. In adult workers, the *lncov1*/Tudor-SN module responds positively to environmental cues that suppress reproductive capacity. As *lncov1* is considerably conserved in the Apidae, we propose that, by promoting worker sterility, the *lncov1*/Tudor-SN module has likely played critical roles in the social evolution of bees.

## INTRODUCTION

Sociality in insects is a major evolutionary transition that has occurred independently in the Hymenoptera (bees, wasps, and ants), and in the cockroach-related termites (Isoptera). The apex of sociality in the Hymenoptera is marked by the presence of morphologically and physiologically distinct female castes, the queen and the worker (Korb and Heinze, 2016; Linksvayer and Johnson, 2019). These are mutually dependent, particularly during the sessile phase of the colony life cycle. This strong interdependence represents an evolutionary point of no return, as it prevents the reversion to a communal or even solitary life history (Boomsma and Gawne, 2018; Wilson and Hölldobler, 2005). Species with such reproductive division of labor are referred to as being ‘eusocial’ (Michener, 1974; Sherman et al., 1995; Thompson and Oldroyd, 2004) and, specifically, as highly eusocial when they have morphologically distinct castes.

There are two major open questions concerning the female caste dimorphism in eusocial insects: (a) how did social insects go beyond phenotypic plasticity to stably generate the two irreversibly distinct castes (Sommer, 2020), and (b), what is the configuration of the regulatory networks that drive the divergent ontogenetic pathways of the two castes (Friedman et al., 2020)? Previous studies have examined a set of bee species that span the full spectrum of social organization from solitary to the highly eusocial (Fischman et al., 2011; Kapheim et al., 2020; Rehan and Toth, 2015). These comparative transcriptomic and genomic analyses have identified gene families and molecular signatures potentially associated with the evolution of eusociality. In addition, epigenetic mechanisms, as key regulators of chromatin activity (Allis and Jenuwein, 2016), have been proposed to play roles in the morphological, physiological and behavioral differentiation between and within genetically identical females (Alvarado et al., 2015; Duncan et al., 2020; Kucharski et al., 2008). Yet, understanding how these genes are functionally integrated into regulatory gene and epigenetic networks for building the alternative caste phenotypes remains a challenging task, even in the model organism for insect sociality, the Western honey bee, *Apis mellifera* L. (Barchuk et al., 2007; Cameron et al., 2013; Evans and Wheeler, 1999; Foret et al., 2012; Wojciechowski et al., 2018). A major hurdle to be overcome in this context is to distinguish between regulatory pathways involved in driving general body growth (queens are larger than workers) from those that coordinate the caste-specific development of tissues and organs, especially in the female reproductive system.

The most remarkable caste difference between honey bee queens and workers lies in the anatomical architecture of their reproductive systems, particularly their ovaries. While each of the paired ovaries of an adult honey bee queen contains on average 150 ovarioles (the filamentous units comprising the insect ovary), those of a typical worker bee contain on average only two to four ovarioles (Leimar et al., 2012). Yet, as young larvae, queens and workers start out equally, with the same number of ovariole primordia. It is only during the transition from the penultimate (fourth) to the last (fifth) larval instar that a massive programmed cell death (PCD) destroys well over 90% of the ovariole primordia in workers (Hartfelder et al., 2018). These PCD events are inhibited by a topical application of synthetic juvenile hormone-III (Schmidt-Capella and Hartfelder, 1998) (JH), an insect hormone that, together with ecdysteroids, orchestrates the postembryonic molts in insects (Bellés, 2020). Elevated JH levels in the hemolymph of young queen honey bee larvae is critical for protecting their ovaries from PCD, resulting in the large ovary phenotype (Rachinsky et al., 1990). Despite PCD events, honey bee workers still retain limited reproductive capacity as adults, which is repressed by pheromone signals released by the queen and her brood (Slessor et al., 2005). Hence, JH biosynthesis genes are implicated in caste-specific developmental trajectories in social bees (Bomtorin et al., 2014; Cardoso-Júnior et al., 2017).

While transcriptomic analyses of honey bee ovaries have provided insights into potential gene regulatory networks (Duncan et al., 2020; Lago et al., 2016), only a few specific genes have actually been functionally characterized thus far. For instance, the functional knockdown of the DNA methyltransferase 3 by RNA interference (RNAi) resulted in a queen-like phenotype, mirroring the dietary effects induced by the royal jelly (Kucharski et al., 2008). Moreover, genes associated with the insulin/insulin-like and target of rapamycin signaling (IIS/TOR) pathways likely play roles in honey bee caste dimorphism as they link nutrient-sensitive pathways to downstream regulators of reproduction (de Azevedo and Hartfelder, 2008; Patel et al., 2007).

A candidate gene of particular interest regulating caste dimorphism is the long non-coding RNA (lncRNA) denominated *long non-coding ovary-1 RNA* (*lncov1*) (Humann and Hartfelder, 2011; Humann et al., 2013). lncRNAs, usually defined as ncRNAs longer than 200 bp (Ma et al., 2013), have been hypothetically implicated in the behavioral plasticity (Glastad et al., 2019; Liu et al., 2019; Yan et al., 2014) and ovary activity (Chen and Shi, 2020; Chen et al., 2017) of social insects. Specifically, *lncov1* is genomically located on chromosome 11, in an intron of a predicted gene of unknown function, designated *LOC726407*. Interestingly, the *LOC726407* gene (and thereby, *lncov1*) maps within a quantitative trait locus (QTL) associated with variation in ovariole number in adult honey bee workers (Humann et al., 2013; Linksvayer et al., 2009). Importantly, *lncov1* is overexpressed in the ovaries of worker larvae just as they undergo PCD, while expressed at basal levels in queen ovaries (Humann and Hartfelder, 2011). Together, this makes *lncov1* a strong candidate for regulating the divergence in reproductive capacity between honey bee queens and workers.

While functions for lncRNAs has so far only been proposed in neuronal processes of adult honey bee workers (Kiya et al., 2012; Sawata et al., 2004) and ants (Shields et al., 2018), here we mechanistically and functionally investigate the *in vivo* roles of *lncov1* in worker sterility of honey bees kept under natural conditions or subjected to endocrinal, social and dietary treatments that modulate ovary activation.

## RESULTS

### The expression of *lncov1* and *LOC726407* are negatively associated across the development of honey bee larvae

To gain insights into the expression dynamics of *lncov1* and its protein coding host gene *LOC726407,* we determined their transcript levels by quantitative PCR (RT-qPCR) in four tissues of the fourth larval instar (L4) and six substages of the fifth larval instar (L5). We found that *lncov1* is higher expressed in the fat body and the ovaries compared to head tissue and the leg imaginal discs (Fig. 1A). We also provide confirmative evidence for its expression peak previously reported in the ovaries late feeding-stage fifth instar worker larvae (L5F3) (Humann et al., 2013). *LOC726407* expression is higher in the head and the leg imaginal discs compared to the two abdominal tissues (Fig. 1A). This indicates that the expression of the intronic *lncov1* transcript is specifically regulated and not a by-product of mRNA processing of its host gene, *LOC726407*. Furthermore, a correlation analysis revealed a negative association between *lncov1* abundance and *LOC726407* gene expression (Fig. 1B, two-tailed Spearman’s correlation test, ρ = -0.768, *p* < 0.0001, n = 82).

**Figure 1.**
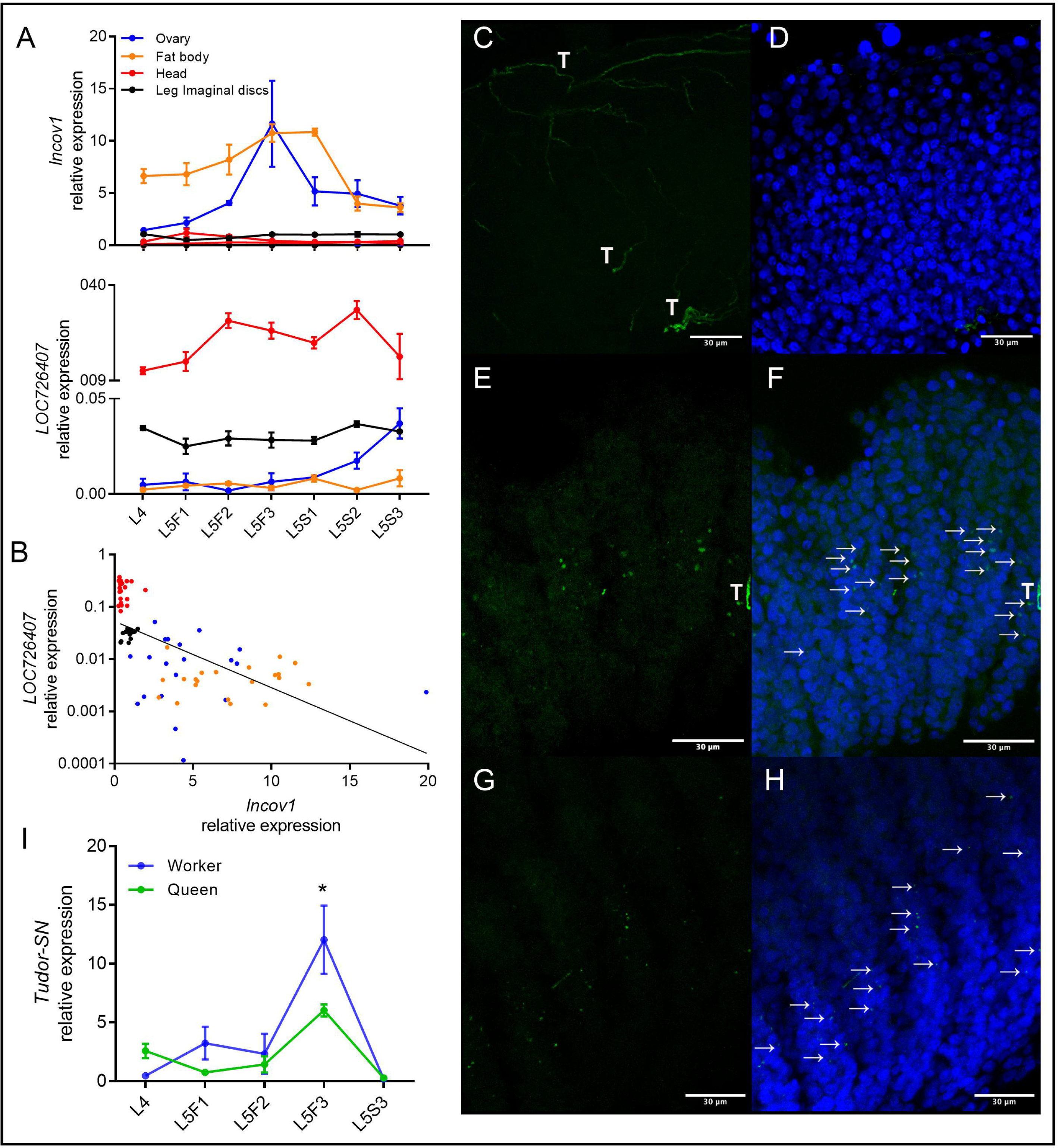
Transcriptional dynamics of *lncov1, LOC726407,* and *Tudor-SN* genes in the larval ovaries of honey bees. **A.** Temporal dynamics of *lncov1* (upper graph) and *LOC726407* (bottom graph) expression in larval workers. Four tissues were investigated: ovaries (blue), head (red), fat body (orange), and leg imaginal discs (black), in seven larval developmental stages: fourth larval stage (L4); and fifth instar feeding and spinning stage larvae (L5F1 – L5S3) [the *lncov1* expression data shown in a for the ovaries is from (Humann et al., 2013)]; shown are the means ± SEM. **B.** *LOC726407* expression plotted against *lncov1* expression across the samples shown in **A**; the black line indicates the trend (Two-tailed Spearman’s rank correlation test, ρ = -0.768, *p* < 0.0001, n = 82). **C-H**. Fluorescence *in situ* hybridization (FISH) of *lncov1* transcripts in the ovaries of larval honey bee workers. **C-D.** L4 stage. **E-F.** L5F3 stage. **G-H.** L5S3 stage. The FISH images on the left (**C, E, G**) show the FISH signal for *lncov1* in green, those on the right (**D, F, H**) show the overlay of the *lncov1* fluorescence (green) in relation to the nuclear stain DAPI (blue); white arrows indicate *lncov1* speckles and T indicates autofluorescent tracheoles present in the ovaries. Scale bars 30 μm. **I**. Temporal dynamics of *Tudor-SN* expression in the ovaries of queen (green) and worker (red) larvae. Shown are the means ± SEM (Two-way ANOVA, \**p* < 0.01, Table S2). See Table S6 for details of sample size.

### The spatial and temporal *lncov1* expression correspond with PCD events in larval worker ovaries

Using fluorescence *in situ* hybridization (FISH) we identified the ovary regions where *lncov1* is expressed (Fig. 1C-H). In fourth instar (L4) worker ovaries, i.e., prior to the onset of PCD, *lncov1* transcripts are not detectable (Fig. 1C-D). However, in L5F3 worker ovaries, at the peak of *lncov1* expression, the transcripts are visible as clear fluorescence signals, especially in the central region of the ovarioles (Figs. 1E-F and Fig. S1A,B). At this developmental stage, strong TUNEL-positive marks indicating PCD have previously been detected exactly in this central region of the ovarioles (Schmidt-Capella and Hartfelder, 1998). The intracellular localization of *lncov1* transcripts in the L5F3 ovarioles appeared as speckles and diffused throughout the cytoplasm of the germline cells (Fig. S1A,B). The cytoplasmic localization of *lncov1* RNA revealed by the FISH assays is corroborated by *in silico* predictions (iLoc-LncRNA software) of its subcellular location.

In L5S3 worker ovaries, the FISH-positive *lncov1* fluorescence appeared to be reduced compared to the L5F3 stage (Fig. 1G-H), consistent with the lower *lncov1* transcript levels in the L5S3 ovaries (Fig. 1A). Furthermore, the *lncov1* speckles were spread out along the entire ovariole axis and no longer concentrated in the central region (Fig. 1G-H). This expression pattern again corresponds with the more widespread TUNEL-positive marks previously found for this developmental stage (Schmidt-Capella and Hartfelder, 1998). In the negative control samples prepared with the sense probe no signal was detected (Fig. S1C-H). These FISH results, not only provide independent support for the temporal dynamics of the *lncov1* transcript levels obtained from the RT-qPCR assays (Fig. 1A), but also establish that its expression peak is spatially associated with the onset of PCD in the ovaries of worker larvae.

### *lncov1* interacts with the cell death-associated Tudor-SN protein

For a deeper understanding of the role of the *lncov1* gene in honey bee development we identified putative interaction partners by performing pull-down assays followed by shotgun proteomics (LC-ESI-MS/MS). For this, the full-length *lncov1* RNA was expressed *in vitro*, immobilized on magnetic beads, and then incubated with protein extracts from L5F3 worker larvae. Mass spectrometry analyses of the pull-down products identified the following *lncov1*-interacting proteins: Tudor-SN, with nine hits in the Mascot database to *A. mellifera* proteins; Elongation factor 1-beta, with three Mascot hits; Elongation factor 1-gamma-like, also with three Mascot hits; Elongation factor 1-delta-X2, with four Mascot hits; 3-ketoacyl-CoA thiolase, mitochondrial isoform X2, with five Mascot hits; and 60S acidic ribosomal protein P1, with one Mascot hit (Table S1).

The protein with the highest Mascot scores, Tudor-SN, is a highly promising candidate to reveal predictive *lncov1*-sterility-associated functions because Tudor-SN is an evolutionarily-conserved component of the apoptosis degradome in plants and animals, essential for PCD propagation (Sundström et al., 2009). In addition, the honey bee Tudor-SN protein is *in silico* predicted to be cytoplasmic (ProtComp v9.0 software), suggesting that the *lncov1*/Tudor-SN complex may be assembled and functional in the cytoplasm of the ovariole cells of honey bee worker larvae. Therefore, the interaction of *lncov1* and Tudor-SN protein provides a plausible functional explanation for the cytoplasmic localization of *lncov1* observed in the FISH assays. Our downstream functional analyses were, therefore, focused on this candidate protein.

### *Tudor-SN* expression coincides with the expression of *lncov1* and is elevated in the ovaries of worker larvae undergoing PCD

As Tudor-SN had not previously been implicated in honey bee worker sterility, we first quantified its expression levels in the ovaries of queen and worker larvae in the critical larval stages (L4 and L5). The RT-qPCR data not only showed that *Tudor-SN* is expressed in the larval ovaries of both queens and workers, but also that it has an expression peak in L5F3-stage worker ovaries (Fig. 1I and Table S2), perfectly coinciding with the *lncov1* expression peak (Fig. 1A). This temporal correlation for the expression of the two genes provides additional evidence for the physical *lncov1*/Tudor-SN interaction shown through the pull-down assay.

### The silencing of *Tudor-SN* affects the expression of pro-apoptotic genes and effector caspase activity

For direct functional insights into the interaction of *lncov1*/Tudor-SN we performed RNAi experiments in honey bee larvae. To do so, *lncov1* or *Tudor-SN* double-strand RNAs (ds-*lncov1* and ds-*Tudor-SN*, respectively) was added to the diet of L5F2-stage worker larvae, i.e., just prior to the expression peaks of both genes. In the ovaries of ds*-Tudor-SN*-treated larvae, the expression of *Tudor-SN* was reduced by approximately 50% compared to untreated larvae or the negative controls treated with *ds-GFP* (Fig. 2A and Table S3). In ds-*lncov1* treated larvae, however, we were unable to achieve a significant knockdown of the target gene, despite several attempts and using both *in vivo* and *in vitro* dsRNA treatment strategies (Fig. S2A-C). In the subsequent experiments we, therefore, focused on the effect of the *Tudor-SN* knockdown on molecular pathways known to be associated with honey bee worker sterility.

**Figure 2.**
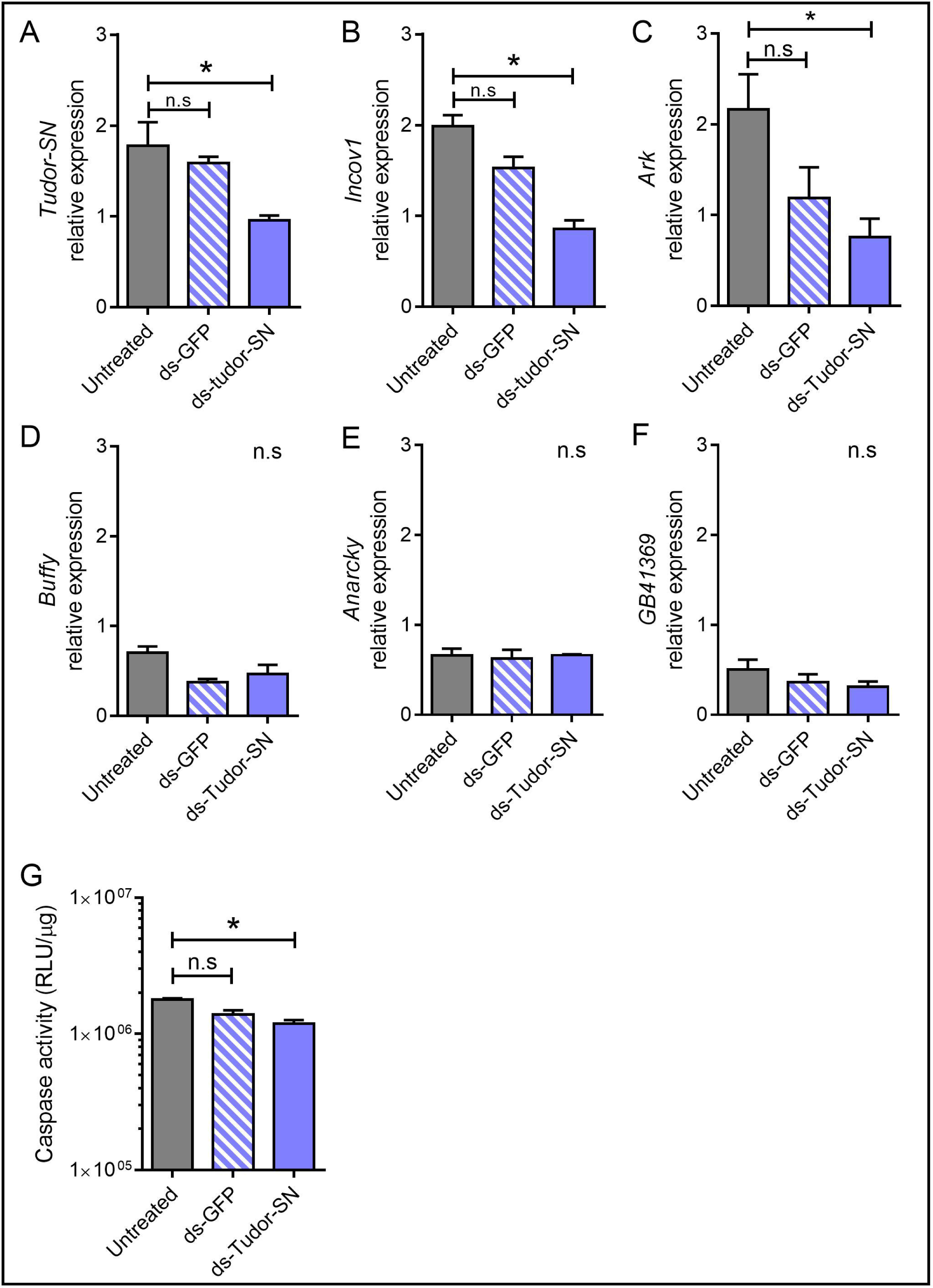
Effect of the RNAi-mediated knockdown of *Tudor-SN* function in the ovaries of worker larvae. Treatment groups: ds-*Tudor-SN*, negative control *ds-GFP,* and untreated larvae. **A.** Knockdown efficiency of ds-*Tudor-SN*. **B.** Relative expression of *lncov1*, *Ark* (**C**), *Buffy* (**D**), *Anarchy* (**E**) and *GB41369* (**F**), a putative effector caspase coding gene. **G.** Effector caspase activity, relative light units (RLU) were normalized to the protein concentration of the sample. Bars represent means ± SEM, \**p* < 0.05 and “n.s” indicates *p* > 0.05. See Table S6 for details of sample size.

We investigated whether the knockdown of *Tudor-SN* affected the expression of its interaction partner *lncov1* and of four worker-sterility related genes: *Anarchy* (Oxley et al., 2008; Ronai et al., 2016a), *Ark* an *Buffy* (Dallacqua and Bitondi, 2014) and the PCD effector caspase *GB41369* (Ueno et al., 2009). We found that worker larvae treated with *ds-Tudor-SN* had significantly reduced expression levels of *lncov1* and *Ark* (Fig. 2B,C and Table S3), while the expression of *Buffy*, *Anarchy* and *GB41369* was unaffected (Fig. 2D-F and Table S3).

Given that the human Tudor-SN ortholog is itself cleaved by the effector caspase Caspase-3 (Sundström et al., 2009), we next conducted a biochemical assay to investigate whether the knockdown of *Tudor-SN* affected effector caspase activity, as a proxy of PCD in honey bees (Ronai et al., 2016b, 2017). Larvae fed with ds*-Tudor-SN* had significantly reduced effector caspase activity compared to control larvae (Fig. 2G and Table S3). In summary, our results show that the knockdown of *Tudor-SN* downregulated the expression of two pro-apoptotic transcripts, *lncov1* and *Ark*, and also modulated the activity of an effector protein of PCD. These results indicate that the *lncov1/*Tudor-SN interaction constitutes a functional module that drives PCD in the ovaries of worker larvae.

### Juvenile hormone treatment does not significantly affect *lncov1 and Tudor-SN* expression

Since queen larvae have hemolymph JH levels that are approximately 10 times higher than worker larvae throughout their larval development (Rachinsky et al., 1990), and the fact that JH inhibits the ovarian PCD (Schmidt-Capella and Hartfelder, 1998), we tested whether JH might be an upstream negative regulator of the *lncov1*/Tudor-SN module. Hence, we applied synthetic JH-III topically to worker larvae, while controls were treated with acetone (solvent control), or left untreated. As expected, the expression of *Kruppel-homolog 1(Kr-h1)*, a primary target of JH in insects (Bellés, 2020), including honey bees (Lago et al., 2016), was significantly up-regulated in response to the JH-III treatment (Fig. S2D-F and Table S4). The mean expression level of *lncov1* was lower in the JH-III treated larvae, but differences were not statistically significant (Fig. S2E and Table S4). The mean expression level of *Tudor-SN* was higher in the JH-III treated larvae, but again, due to the large between-sample variation, this result was also statistically not significant (Fig. S2F and Table S4). Although these expression trends were in the direction expected if JH plays a suppressive role in the *lncov1*/*Tudor-SN* regulatory module, our analysis does not distinguish whether JH signaling and the *lncov1*/Tudor-SN module are in the same or in parallel pathways that modulate PCD events in the caste-specific development of honey bee larval ovaries.

### Queen presence and diet modulate *lncov1*/Tudor-SN activity in adult workers

After finding evidence that the *lncov1*/Tudor-SN module acts as a regulator of PCD activity in the ovaries of honey bee larvae, we next examined whether this module might also be associated with ovary sterility in adult workers. We particularly investigated whether the expression of *lncov1* and *Tudor-SN* is affected by environmental cues known to regulate ovary activation in adult workers (Altaye et al., 2010; Cardoso-Júnior et al., 2021a; Lin and Winston, 1998; Ronai et al., 2016b), such as the presence or absence of a queen, or a nutrient-enriched diet.

We first mimicked the presence/absence of a queen by housing young adult workers in cages containing synthetic queen mandibular gland pheromone (QMP). We found that exposure to QMP led to a significant increase in *lncov1* expression in the ovaries of 4-day-old adult workers, while *Tudor-SN* expression was decreased (Figure 3A and Table S5). Furthermore, the expression of these two genes changed according to the age of the adult workers.

**Figure 3.**
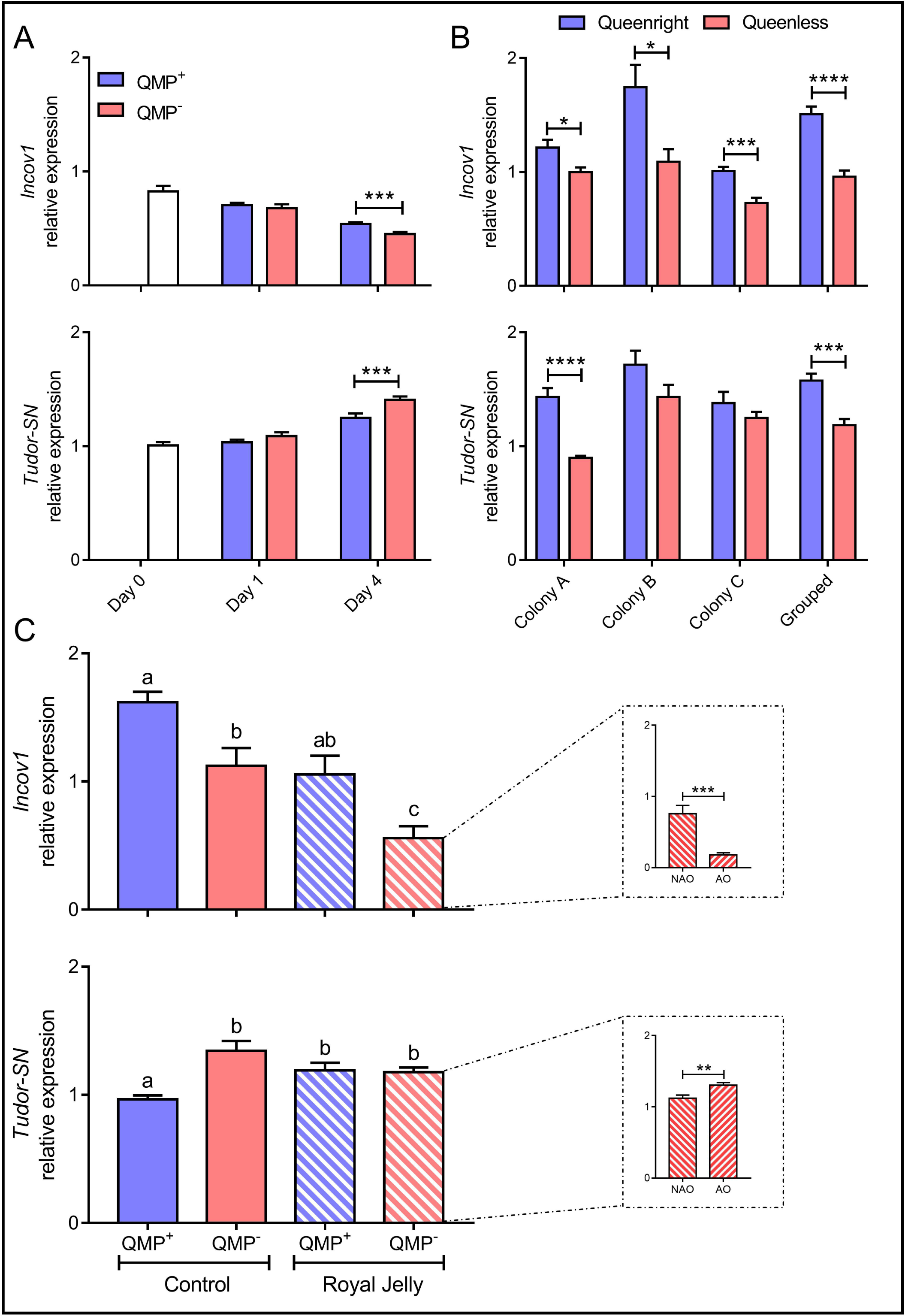
Modulation of the expression of *lncov1* and *Tudor-SN* in the ovaries of adult workers by the presence of a queen and diet. **A.** 1- and 4-day-old caged workers exposed (QMP^+^, blue bars) or not exposed (QMP^-^, red bars) to a strip of synthetic queen mandibular pheromone (QMP). Workers collected before treatment (Day 0) were used to determine the expression baseline. Shown are means of relative expression ± SEM for *lncov1* (upper graph) and *Tudor-SN* (bottom graph), \*\*\**p* < 0.001. **B.** 4-day-old workers kept in queenright (QR, blue bars) or queenless (QL, red bars) field colonies. Shown are means of relative expression ± SEM for *lncov1* (upper graph) and *Tudor-SN* (bottom graph), \**p* < 0.05, \*\*\**p* < 0.001, \*\*\*\**p* < 0.0001. **C.** Combined effects of diet and social environment on *lncov1* (upper graph) and *Tudor-SN* (bottom graph) expression. Workers were fed a control diet (full bars) or a royal-jelly-rich diet (striped bars) for seven days, while they were at the same time exposed (QMP^+^, blue bars) or not to synthetic QMP (QMP^-^, red bars). For the QMP^-^/royal jelly bees, the expression of *lncov1* and *Tudor-SN* were evaluated separately for workers with non-activated (NAO) and activated ovaries (AO) (insets: Student’s t-test, \*\**p* < 0.01, *** *p* < 0.001). Bars represent the means ± SEM; different letters indicate a statistical difference at *p* < 0.05 for GLMM tests. See Table S6 for details of sample size.

Since experiments with caged bees can only partially reflect what happens in a real colony, we next quantified the levels of *lncov1* and *Tudor-SN* expression in the ovaries of 4-day-old sister workers of paired queenright or queenless colonies kept under field conditions. As with the results for the caged bees, we found that *lncov1* expression is significantly higher in the workers of all three queenright colonies relative to their queenless pair (Figure 3B and Table S5). The results for *Tudor-SN* expression, however, were significantly higher in one queenright colony (colony A), and showed no difference for the other two colonies (Figure 3B and Table S5). When the data from all three colonies are pooled, *Tudor-SN* expression is significantly higher in the queenright condition (Fig. 3B and Table S5), in contrast to what is found for the cage experiment. This effect was not entirely driven by colony A, as the trend in the other two colonies appeared to be in the same direction. In conclusion, *lncov1* is overexpressed in response to a queen signal, either in the cage or field colonies; however, we found an effect of the queen on *Tudor-SN* expression in controlled cage experiments but this finding is in the opposite direction for the field colony study.

Second, we investigated whether the adult workers’ diet affects the *lncov1*/Tudor-SN module. For this we fed newly emerged workers a queen-like diet consisting of highly nutritious royal jelly (RJ) mixed with honey to induce ovary activation (Cardoso-Júnior et al., 2021a), or a worker (control) diet, consisting of 50% honey. Workers kept in RJ or control cages, exposed or not to QMP, were provided pollen (protein source) and water *ab libitum*.

In accordance with the hypothesis that *lncov1* suppresses ovary activation in adult bees, we found that workers fed RJ-rich diet significantly reduced *lncov1* transcription in their ovaries, while QMP upregulated *lncov1* levels (Fig. 3C and Table S5). Furthermore, the expression pattern of *lncov1* was exactly opposite to the ovary activation score (Cardoso-Júnior et al., 2021a), indicating that *lncov1* expression is negatively associated with ovary activation in adult worker (Fig. S3, Two-tailed Spearman’s correlation test, ρ = -0.6, *p* < 0.0001, n = 76). To test for a direct role of *lncov1* in ovary activation in adult workers, we quantified the *lncov1* expression levels in activated and non-activated ovaries of QMP^-^/RJ workers as this group had a sufficient number bees with activated ovaries that it could be analyzed separately. We found that the expression of *lncov1* was significantly lower in the activated ovaries compared to non-activated ones (Fig. 3C inset and Table S5). On the other hand, *Tudor-SN* expression is enhanced by the RJ diet but suppressed by QMP (Fig. 3C and Table S5). Furthermore, *Tudor-SN* expression is higher in activated ovaries compared to non-activated ones (Fig. 3C inset), and positively correlated with the ovary activation score (Two-tailed Spearman correlation test, ρ = 0.31, n = 76, *p* = 0.006).

Thus, our results indicate that the queen, via her mandibular gland pheromone, triggers a consistent up-regulation of *lncov1* expression in adult worker bees as a component of the pathway that suppresses their ovarian activation. In contrast, the RJ-enriched diet attenuated the inhibitory effect of QMP on the ovarian *Tudor-SN* expression level (Fig. 3A,C) and, in a more general sense, ovary activation (Cardoso-Júnior et al., 2021a).

### Independent validations of *lncov1*’s roles in honey bee reproduction

To independently validate our findings that all point towards a central role of *lncov1* in honey bee worker sterility, we investigated whether and to what degree *lncov1* and *Tudor-SN* appear as differentially expressed in publicly available RNA-Seq datasets (Chen and Shi, 2020; Chen et al., 2017; Duncan et al., 2020). These transcriptomes were downloaded and reanalyzed with a focus on the expression of three genes of interest, *lncov1*, *Tudor-SN*, *LOC726407. Gadph* (GB50902) expression levels were checked as endogenous control for between-sample variability. These transcriptomes permitted us to independently assess the expression of these genes in activated ovaries from queenless workers and queens in comparison to the inactivated ovaries of queenright workers.

For the RNA-seq libraries generated by Duncan and colleagues (Duncan et al., 2020), *lncov1* is overexpressed in the inactivated ovaries of queenright workers, while reduced transcript levels is found in the ovaries of queenless workers and queens (Fig. 4A). Contrary to the clear pattern for *lncov1*, the expression of *Tudor-SN* does not differ between activated and inactive ovaries (Fig. 4B), supporting our own results for colony-reared bees (Fig. 3B). The expression of *LOC726407* followed the same patterns of *lncov1* (Fig. S4), indicating that the negative association between *lncov1* and its host gene *LOC726407* (Fig. 1B) might be restricted to larval development and/or is more evident in other tissues of adult bees. As there was no difference with respect to *Gapdh* expression (Fig. S4), the differences observed for the *lncov1* and *LOC726407* transcripts likely reflect real biological differences between the two ovary types and are not due to technical issues or between-sample variability.

**Figure 4.**
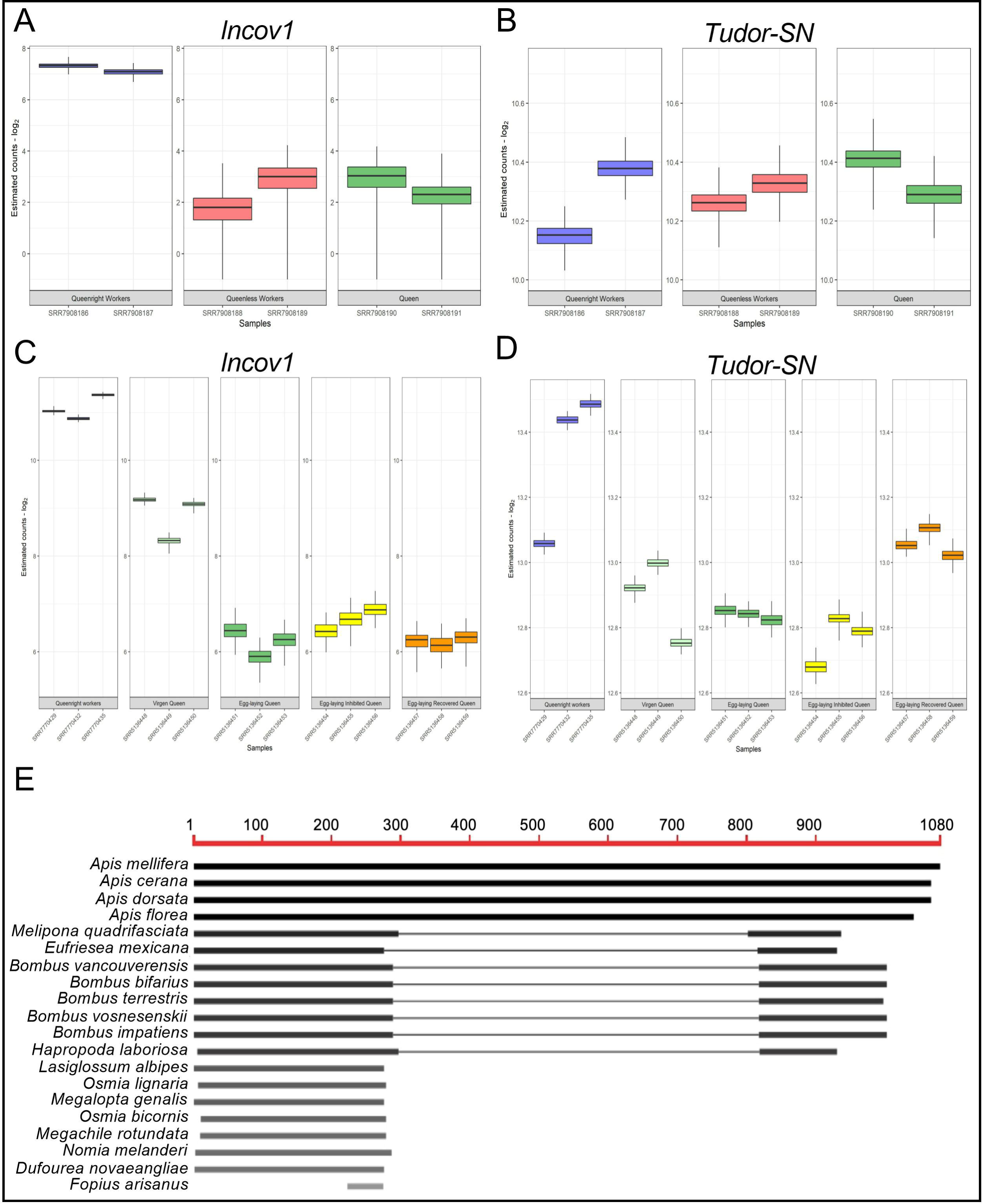
Alignments of the *A. mellifera lncov1* gene in the genomes of other social insects, and quantification of *lncov1* and *Tudor-SN* transcripts in published RNA-seq datasets. **A**. Estimated read counts for *lncov1* and *Tudor-SN* (**B**) transcripts in the ovary of queenright workers (blue), queenless workers (red) and queen (green) found in the RNA-seq libraries published by Duncan et al., 2020. Boxplots represent the intern quartiles of 1000x bootstrap independent read counts. **C**. Estimated read counts for *lncov1* and Tudor-SN (**D**) transcripts in the ovary of queenright workers (blue), virgin queen (light green), egg-laying queens (green), caged queens (“egg-laying inhibited queen” - yellow) or caged-released queens (“egg-laying recovered queen” – orange) found in the RNA-seq libraries published in the “Chen and Shi, 2020” and “Chen et al., 2017” studies. Boxplots represent the inner quartiles after 1000x independent bootstrap read counts. **E.** *lncov1* blast against the genomes deposited in HymenopteraMine. Except for *F. arianus*, which is a braconid wasp, all others are bees. The grey scale level of the bars indicates the degree of similarity with *A. mellifera lncov1.* Figure obtained from HymenopteraMine. For details on sequence IDs and alignment scores, see Table S8.

We also investigated the expression of the same four genes in the ovarian transcriptomes of workers and adult queens recently published by the same research group (Chen and Shi, 2020; Chen et al., 2017). *lncov1* is overexpressed in the ovaries of adult queenright workers, while moderate to low expression levels are found in the ovaries of virgin queens and mated queens, respectively (Fig. 4C). Ovarian expression of *Tudor-SN* is high in workers and shows medium-low expression levels in the ovaries of queens (Fig. 4D). Interestingly, *Tudor-SN* expression is increased in all three samples of queens that had been allowed to re-start oviposition when compared to other mated queens (Fig. 4D). This pattern in *Tudor-SN* expression was not observed for the other genes assessed (Fig. 4C and S6). Taken together, these results provide compelling independent evidences for the hypothesis that the *lncov1*/Tudor-SN regulatory module is indeed key to the flexible modulation of the reproductive activity of honey bee queens and workers.

Since this analysis revealed that *lncov1*, as shown by the logarithmic scale in Fig. 4A,C, is overexpressed in the ovaries of queenright workers, we determined whether *lncov1* would appear in the list of differentially expressed genes considering whole ovary transcriptome. Hence, we computationally included the *lncov1* sequence as a gene ID to the list of transcripts generated by the honey bee genome, and after FDR corrections, we found that *lncov1* is listed within the differentially expressed transcripts in all the scenarios tested (Fig. S5). Together with our RT-qPCR data, these *in-silico* analyses are independently generated evidences in favor of the hypothesis that elevated levels of *lncov1* in the ovaries of queenright workers are an intrinsic signature of worker sterility.

### *lncov1* is an evolutionarily conserved lncRNA in Apidae

Next, we investigated whether and to what extent *lncov1*/*Tudor-SN* may be evolutionarily conserved in the genomes of bees and hymenopterans in general. Using the *A. mellifera lncov1* sequence as query for blastn searches for species in the Hymenopteran Genome Database (Elsik et al., 2018), we found that *A. mellifera lncov1* has a nearly full-length alignment (E-value 0.00) with sequences of the three other honey bee species (genus *Apis*): *A. cerana, A. dorsata,* and *A. florea* (Fig. 4E and Table S8). High levels of sequence conservation is also seen for the other three branches of corbiculate bees, represented by the orchid bee *Eufriesea mexicana* (solitary to facultative eusocial), the stingless bee *Melipona quadrifasciata* (highly eusocial), and five bumble bee species (genus *Bombus;* primitively eusocial). In these non-*Apis* corbiculates, as well as in the anthophorid *Habropoda laboriosa,* we found two highly conserved fragments of the *lncov1-*like gene, one of 290 to 298 bp corresponding to the 5’end of *A. mellifera lncov1* and a shorter one of 121 to 190 bp fragment with significant similarity to the 3’end of *A. mellifera lncov1* (Fig. 4E and Table S8). In *Ceratina calcarata*, a member of the family Apidae, tribe Ceratini, two structural elements of *A. mellifera lncov1* were also identified, but with much lower levels of similarity (Table S8). In the genomes of five solitary bees, all belonging to the families Halictidae and Megachilidae, sequence conservation was restricted to the 281 to 297 bp fragment of the 5’end of *A. mellifera lncov1.* Outside the bees (Anthophila), only a small fragment of 51–53 bp in length located at the end of this conserved 5’ region showed significant similarity (e^-9^ to e^-6^) with honey bee *lncov1* (Fig. 4E and Table S8). This fragment was found in the genomes of ants (Formicidae) and parasitic wasps (Braconidae). Thus, we found remarkably high sequence conservation for honey bee *lncov1* in the other species of the genus *Apis*. Considerable sequence conservation was also noted in the other social corbiculate bees and, gradually decreasing in the other bee families whose females generally have a solitary lifestyle. This finding provides additional support for the hypothesis that *lncov1* played a critical role in the social evolution of bees.

## DISCUSSION

We functionally characterized the first lncRNA implicated in the evolution of highly eusocial insects, the *lncov1*. Its expression patterns in the ovaries of honey bee larvae, together with its responsiveness to environmental cues that modulate ovary activity in adults, strongly suggest *lncov1* is associated with the ovarian PCD events that establish and maintain the worker ovaries in a quiescent status. The *lncov1*-interaction partner, the Tudor-SN protein, is an essential component of the PCD degradome from animals to plants serving as a substrate for effector caspase cleavage during apoptosis propagation (Sundström et al., 2009). Moreover, Tudor-SN wild-type functions are essential for cell viability as it participates in microRNA biogenesis mechanism being part of the RNA-induced silencing complex (Caudy et al., 2003). Based on these findings, we propose a model where *lncov1* is a key component of the molecular machinery that drives honey bee worker sterility through its interaction with Tudor-SN (Fig. 5), a regulatory mechanism likely generalized to other social bee species.

**Figure 5.**
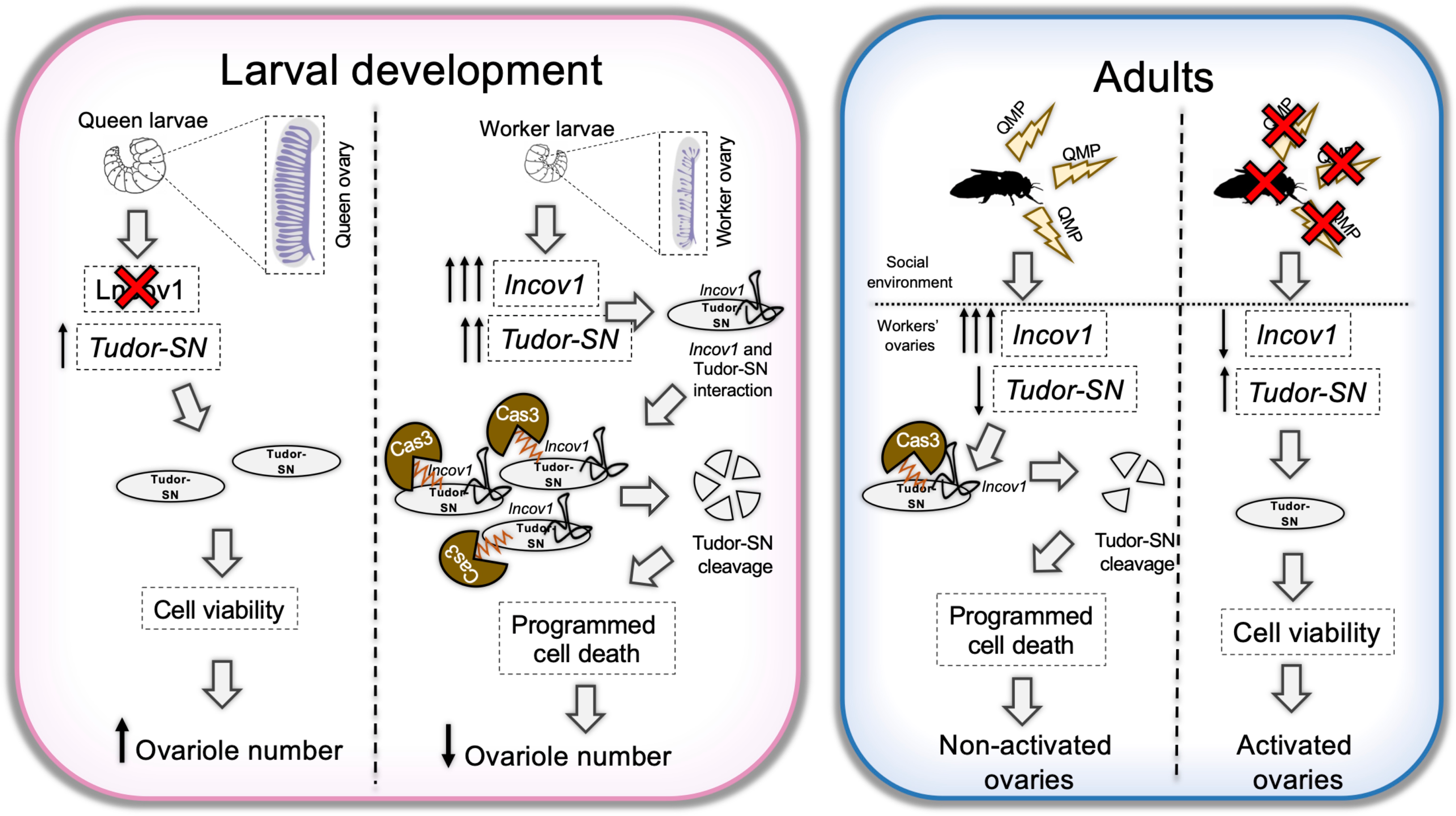
Model of the *lncov1*/Tudor-SN regulatory mechanisms triggering PCD that promote worker sterility in honey bees. Left panel (pink): As a result of differential feeding regimes (i.e., royal jelly *vs*. worker jelly), worker larvae express high levels of *lncov1* and *Tudor-SN* in their ovaries. In contrast, the ovaries of queen larvae express basal levels of *lncov1* (Humann and Hartfelder, 2011 and Peruzzollo et al. unpublished results), while retain moderate levels of *Tudor-SN* expression. In the absence of *lncov1*, Tudor-SN protein is not degraded in the ovaries of queen larvae and can act in the RNA-induced silencing complex. In contrast, when the *lncov1*/Tudor-SN regulatory module is assembled in the ovaries of worker larvae, it likely triggers the degradation of Tudor-SN by caspase effector proteins, thus propagating apoptosis. Right panel (blue): In adult honey bee worker ovaries, *lncov1* is overexpressed in response to queen signals and a worker-like poor nutritious diet. Hence, high levels of *lncov1* compromise Tudor-SN functions, resulting in inactive ovaries. However, when a colony lacks a queen, or when a highly nutritious queen-like diet is provided, the workers’ ovaries can become activated decreasing *lncov1* and increasing *Tudor-SN* expressions. This model emphasizes the effect of *lncov1* flexibly adjusting ovariole status between death and viability, a process that operates throughout honey bee life cycle.

We suggest that Tudor-SN has a Janus function that depends on the presence or absence of *lncov1.* In the presence of *lncov1,* as is the case in larval and adult queenright worker ovaries, Tudor-SN is required to propagate PCD events, whereas in the absence of *lncov1* (queen ovaries and ovaries of queenless workers), Tudor-SN likely promotes cell viability. Supporting this model, the Tudor-SN protein was first characterized in *D. melanogaster*, where a loss-of-function mutation in the *Tudor-SN* sequence caused sterility (Boswell and Mahowald, 1985), and the RNAi-mediated knockdown of a Tudor-SN protein family member in the oriental fruit fly *Bactrocera dorsalis* caused a disruption in ovary development (Xie et al., 2019). Moreover, we found that *lncov1* and *Tudor-SN* have a precise temporal overlap in their expression, suggesting a shared pathway for transcriptional regulation. We confirmed this via an RNAi experiment, where the knockdown of *Tudor-SN* led to a significant reduction in the expression of *lncov1*. Unfortunately, we were not able to test for a reciprocal effect, as we did not obtain a significant RNAi-mediated knockdown when targeting *lncov1*. Nonetheless, Tudor-SN positively regulates the expression of the pro-apoptotic gene *Ark,* which has previously been implicated in PCD in the ovaries of larval honey bees (Dallacqua and Bitondi, 2014). More importantly, the knockdown of *Tudor-SN* lowers the activity of effector caspases at the protein level, indicating that Tudor-SN is itself a substrate of caspase activity in honey bees (this study) as well as in other metazoans (Sundström et al., 2009). Furthermore, our results also indicate that Tudor-SN functions in a PCD pathway that does not involve the anti-apoptotic gene *Buffy*, which is thought to protect the larval ovaries from undergoing PCD (Dallacqua and Bitondi, 2014), or the *Anarchy* gene, which regulates ovary activation in adult honey bee workers (Ronai et al., 2016a). Together with the important evidence that the abortive ovaries of worker larvae express higher levels of *Tudor-SN* than the well-developed ovaries of queen larvae, these results indicates that during the evolution of a worker caste Tudor-SN functions in cell-viability has been co-opted toward its evolutionary-conserved PCD-related roles.

In adult workers then, the *lncov1/*Tudor-SN complex prevents workers from activating their ovaries in the presence of a queen signal, and thus plays key roles in preventing eventual reproductive social conflicts in the colony. Potentially, this PCD pathway acts through the direct activation of *Ark* and effector caspase activity, just as in the larval ovaries, or through interactions with other worker sterility genes such as *Anarchy* (Ronai et al., 2016a, 2017) and Notch signaling (Duncan et al., 2016). These downstream effects are especially interesting in view of the contrasting results between the cage and the colony experiments for *Tudor-SN* expression. It is plausive that other social cues, such as brood pheromones (Slessor et al., 2005; Traynor et al., 2014) might have a role counteracting the effects of the QMP on Tudor-SN expression in the ovaries of workers. For instance, increased levels of Tudor-SN in colony-reared workers is actually predicted to result in a stronger activation of the *lncov1*/Tudor-SN PCD-associated pathway in queenright social context because *lncov1* transcription is active. Thus, a plausive hypothesis to these apparently contrasting results might be that a queen signal unequivocally ensures *lncov1* overexpression while her brood evolved fine-tuning the activation of the *lncov1*/Tudor-SN regulatory complex. We propose that worker ovarian cell death is effectively achieved primarily through overexpressing *lncov1* in response to the presence of a queen, and that this epigenetic factor is sufficient for interrupting Tudor-SN function regardless of transitory changes in *Tudor-SN* expression levels in response to brood or queen pheromones. These results indicate that *lncov1* and its interaction partner Tudor-SN constitute a molecular signature of worker sterility that operates during the entire life cycle of honey bees favoring colony cohesion.

Supportive of a general role of *lncov1*/Tudor-SN operating in the reproduction of both castes, we found changes in the expression of Tudor-SN in the ovaries of honey bee queens when they need to re-start oviposition, a process naturally faced by queens after swarming. Recently, we showed that *lncov1* and *Tudor-SN* expressions are both associated with a reduction in reproductive capacity of adult queens as a result of caging (Aamidor et al., 2022). Together with the findings that *Tudor-SN* is required in the ovaries of queens, both at larval and adult stages, these evidences suggest that the Tudor-SN protein might plays pivotal roles in the reproductive plasticity of queens. To the best of our knowledge, the *lncov1*, through its association with Tudor-SN, is the first lncRNA functionally implicated in the reproductive division of labor in social insects.

A still-open question concerns the function of *lncov1*’s host gene, *LOC726407*. The negative correlation that we found in larvae tissues for the expression of the lncRNA and *LOC726407* suggests that *lncov1* may actually be a negative regulator in *cis* for *LOC726407*. The presence of a CUB (C1r/C1s, Uegf, Bmp1) domain in the amino acid sequence of the LOC726407 protein is interesting because this domain is present almost exclusively in extracellular and plasma membrane-associated proteins that are involved in a wide range of biological functions, including developmental patterning, cell signaling, and fertilization (Blanc et al., 2007). Therefore, it is possible that the low ovarian expression levels of *LOC726407* in workers can be important for reducing their reproductive capacity. Such an inhibitory function of *lncov1* on *LOC726407* expression, however, does not seem to be a general rule, as evidenced by the re-analyses of the transcriptomes of adult honey bees and the positive correlation during brain morphogenesis program of pupal honey bee queens and workers (de Paula Junior et al., 2020). Thus, it is possible that *lncov1* might have pleiotropic functions beyond reproduction, which will likely depend on its interacting partners, time-dependent and and tissue-specific expression profile.

Our finding of the high sequence conservation of *lncov1* in the honey bees (*Apis* genus) is remarkable. Despite knowing that lncRNAs are much less conserved than protein coding genes (Hezroni et al., 2015; Lopez-Ezquerra et al., 2017), especially intronic lncRNAs (Chodroff et al., 2010), sequence similarity of *lncov1* across the genus *Apis* is actually higher than for many protein coding genes. Furthermore, we suggest that *lncov1* is a lineage-specific lncRNA gene that likely originated during the early evolution of bees (Antophila) and became particularly important in the corbiculate bees, a monophyletic clade that originated approximately 80 million years ago (Branstetter et al., 2017; Cardinal and Danforth, 2011). The corbiculate bees comprise the facultatively eusocial orchid bees, the primitively eusocial bumble bees, and the two highly eusocial bees, the honey bees (Apini) and the stingless bees (Meliponini). Recently, *Tudor-SN* was found to be under positive selection in the *Apis* branch when compared to *Bombus* (Fouks et al., 2021). The high sequence conservation of *lncov1* may, thus, have been important for establishing its functional association with the Tudor-SN protein as a regulatory module in the social evolution of bees. In the early stages of sociality, the *lncov1*/Tudor-SN module may have been involved in promoting the reproductive dominance of mothers over their daughters via pheromonal signals and diet (Oi et al., 2015; Ronai et al., 2016c). In the highly eusocial honey bees, however, the *lncov1*/Tudor-SN module may then have been co-opted to promote extensive PCD in the ovaries of worker-destined larvae, defining the reduced number of ovarioles during larval development and ensuring ovary inactivity during adult life cycle. The *lncov1*/Tudor-SN module is, thus, likely a key factor in the evolution of the extreme dimorphism in reproductive capacity of queens and workers that is a hallmark for all the species comprising the genus *Apis*.

The discovery of a novel, functional and active epigenetic pathway that consists of a socially-responsive lncRNA and effector proteins provides insights on the molecular mechanisms underlying the evolution of eusociality in insects. If appreciated under the light of the mammalian epigenetic literature, the findings reported here on the *lncov1*/Tudor-SN module in an invertebrate species are reminiscent of the discovery of Xist, one of the first lncRNAs to be functionally characterized in the early 1990s that is responsible for the X-chromosome inactivation in female mammals (Loda and Heard, 2019). Recently, another “*social*” lncRNA was identified in rodents (Ma et al., 2020), which together with *lncov1*, represents a strong argument that this class of ncRNAs might be of major importance to sociobiology and evolutionary biology. The sophisticated cross-talk between an epigenetic driver of worker sterility (*lncov1*) and its effector proteins further highlights social insects as valuable organismic models to gain evolutionary insights into the epigenetic mechanisms required to express some of the most exuberant forms of polyphenism found in nature, such as the female castes of social insects.

## ACKNOWLEDGMENTS

The authors thank Carlos Couto for his help in the cloning experiments. This work was supported by grants from Funda ão de Amparo à Pesquisa do Estado de São Paulo (FAPESP grants 2016/15881-0, 2017/09269-3 and 2020/08524-2 to CAM, 2011/15810-2 and2014/05757-5 to GJT, 2016/16622-9 to DCL and 2017/09128-0 to KH), the Coordenação de Aperfeiçoamento de Pessoal de Nível Superior - Brasil (CAPES) - Finance Code 001, and the Australian Research Council (DP180101696 to BPO and Amro Zayed).

## SUPPLEMENTARY FIGURE LEGENDS

**Figure S1.** Fluorescence *in situ* hybridization of *lncov1* transcripts in the ovaries of larval workers. **A-B.** Higher magnification of L5F3 worker ovaries showing the cytoplasmic *lncov1* localization as speckles. White arrows indicate *lncov1* speckles (green) and T labels autofluorescent tracheoles present in the ovaries. Nuclei were stained with DAPI (blue). Scale bars 15 μm. **C-H.** Control experiment for the FISH assays using the sense probe for *lncov1* transcripts. The sense probe (control) did not hybridize to *lncov1* transcripts in the ovaries of L4 worker larvae **(C, D)** and neither to those in ovaries of L5F3 **(E, F),** or L5S3 5F3 worker larvae (**G, H).** T marks autofluorescent tracheoles present in the ovaries. Scale bars 50 μm.

**Figure S2.** *In vivo* and *in vitro* RNA interference experiments of *lncov1* in the ovaries of honey bee worker larvae and Juvenile hormone (JH) treatment. **A.** Relative expression of *lncov1* in workers fed double-stranded RNA of *lncov1* (ds-*lncov1*) or *GFP* (ds-*GFP*), or left untreated. **B.** Relative expression of *lncov1* in ovaries cultured *in vitro* in the presence of *lncov1* double-strand RNA (1 µg, 0.1 µg, or 0.01 µg), ds-GFP (1 µg), or left untreated. For all graphs, bars represent relative expression ± SEM (*p* > 0.05). **D.** The relative expression of *Krüppel-homolog 1* (*Kr-h1*), a primary target of the JH response, *lncov1* (**E**) and *Tudor-SN* (**F**) was quantified by RT-qPCR in response to JH-III treatment, acetone (solvent) or left untreated (* indicates statistical significance at *p* < 0.05). For sample size see Table S6.

**Figure S3.** Correlation analysis between the ovary activation score reported in (Cardoso-Júnior et al., 2021a) and the gene expression of *lncov1* (left) and *Tudor-SN* (right). The ovary score of individual samples (gray dots) was determined by averaging the ovary score of all four ovary pairs that compose each sample. The black dashed lines represent the respective trend. Statistical information for two-tailed Spearman’s correlation tests: *lncov1* ρ = -0.601, *p* < 0.0001, n = 76; *Tudor-SN*ρ = 0.312, *p* = 0.006, n = 76).

**Figure S4.** Ovarian expression of *LOC726407* and *Gapdh* transcripts in RNAseq libraries published in (Chen and Shi, 2020; Chen et al., 2017; Duncan et al., 2020). (**A**) Normalized estimated read counts for *LOC726407* and Gapdh (**B**) transcripts in the transcriptomes published in the (Duncan et al., 2020) study. Groups analyzed: queenright workers (blue), queenless workers (red), and queens (green). Boxplots represent the inner quartiles of 1000x bootstrap independent read counts. (**C**) Normalized estimated read counts for *LOC726407* and *Gapdh* (**D**) transcripts in the transcriptomes published in the (Chen and Shi, 2020; Chen et al., 2017) studies. Groups analyzed: queenright workers (blue), virgin queens (light green), egg-laying queens (green), egg-laying inhibited queens (yellow), egg-laying recovered queens (orange).

**Figure S5.** Transcriptomic analyses contrasting the ovarian expression of queenright workers between queenless workers and queens. (**A**) Differentially expressed genes from (Duncan et al., 2020) study are reported in gray dots after FDR correction (adjusted *p* < 0.05, log_2_fold change > 0.5), while red dots represent not significantly differentially expressed genes. Groups being pair-wise compared are reported inside volcano plots (e.g., left side represents upregulated genes in queenright workers, while right side represent genes upregulated in the contrasted group). (**B**) Same as in **A**, but comparing the ovarian expression of queens and workers from (Chen and Shi, 2020; Chen et al., 2017) studies. Groups were: Queenright worker (blue), Virgin queens (light green), Egg-laying queens (“normal queens”, green), Egg-laying inhibited queens (“caged queens”, yellow), Egg-laying recovered queens (“cage released queens”, orange).

## MATERIAL AND METHODS

### Animals and ethics statement

Experiments with honey bee larvae were conducted in Brazil between 2011 to 2017, using Africanized *Apis mellifera* hybrids. Worker larvae were collected directly from brood combs kept in the apiary of the Department of Genetics of the University of São Paulo, Ribeirão Preto, Brazil. All the tissues used in this work were dissected from fourth (L4) and/or fifth (L5) instar larvae staged according to (Rachinsky et al., 1990). The fifth larval instar is subdivided into six substages, these being the F1, F2 and F3 stages when larvae are still feeding, and the S1, S2 and S3 stages, when they have stopped feeding and prepare for metamorphosis.

Experiments with adult honey bees were conducted in Australia between 2017 and 2019, using *A. mellifera ligustica* hives maintained in the apiary of the University of Sydney. Newly-emerged honey bee workers were obtained by keeping sealed brood frames in an incubator at 34.5 °C overnight. For all experiments with adults, each sample consisted of a pool of four pairs of non-activated ovaries, or a single pair of activated ovaries (see Table S6 for sample size details).

Experiments were conducted to reduce animal pain, however, insects, including honey bees, are not subjected to ethical committee approval.

### Gene expression quantification in honey bee larvae

RNA from larvae was extracted from four tissues (ovaries, head, fat body and leg imaginal discs) using TRIzol reagent (Thermo Fisher Scientific, Waltham, MA) following the manufacturer’s protocol, followed by treatment with 0.1 U RNase-free DNase I (Thermo Fisher Scientific, Waltham, MA); see Table S6 for details on sample sizes. RNA quality and quantity were assessed by spectrometry in a Nanovue system (GE Healthcare, Chicago, IL). First strand cDNA was produced using 1 μg of RNA, with oligo(dT) primers and the SuperScript™ II Kit (Thermo Fisher Scientific, Waltham, MA), following the manufacturer’s protocol.

Relative expression was determined for the following genes: *lncov1*, *Tudor-SN*, *LOC726407, Anarchy, Buffy, Ark, Krüppel homolog-1* and *GB41369.* Quantitative RT-PCR (RT-qPCR) analyses were set up using 1 μL of cDNA (diluted 1:10), 7 μL of Power SYBR PCR Green Master Mix (Thermo Fisher Scientific, Waltham, MA), 5 pmol of each primer to a final volume of 14 μL. Reactions were run in triplicate on a Real-Time PCR StepOne Plus system (Life Technologies, Carlsbad, CA) under the following conditions: 50 °C for 2 min, 95 °C for 10 min, 40 cycles of 95 °C for 15 s, and 60 °C for 1 min, followed by melting curve analysis to confirm product specificity. Target gene expression was normalized against *Rp49* (also named *Rpl32*) and *Actin*, which have both been validated as suitable endogenous control genes in honey bee RT-qPCR assays (Lourenço et al., 2008).

For each developmental stage and tissue at least three biological replicates were run (see Table S6). Relative expression was assessed via the 2^-ΔΔCt^ method in both larval and adult samples (see below). Primer sequences are listed in Table S7.

### Expression of *lncov1* and *Tudor-SN* in the ovaries of adult honey bee workers in colonies and cage experiments

The ovaries of adult bees were first macerated in TRIzol reagent (Thermo Fisher Scientific, Waltham, MA), and RNA was then extracted using the Direct-zol RNA MiniPrep Kit (Zymo Research, Irvine, CA). RNA concentrations were determined with a Qubit 2.0 fluorometer (Invitrogen). cDNA was synthesized from 115 ng of RNA using a SuperScript III Reverse Transcriptase Kit (Thermo Fisher Scientific, Waltham, MA) with oligo(dT) primer. The cDNAs were diluted to 2 ng/μL with ultrapure water, and RT-qPCR assays were set up with 2.5 μL SsoAdvanced Universal SYBR Green Supermix (BioRad, Hercules, CA), 1.25 pmol of each primer, 1 μL of diluted cDNA (2 ng), in a total volume of 5 μL. The assays were performed in a CFX384 Real-Time System (Bio-Rad). Three technical replicates were run per sample. RT-qPCR cycle conditions were as follows: 95 °C for 10 min, 40 cycles of 95 °C for 10 s, 60 °C for 10 s, and 72 °C for 15 s, followed by a melting curve analysis. The expression of *lncov1* and *Tudor-SN* was normalized to two reference genes (*Rpl32* and *ef1α*) (Lourenço et al., 2008). The expression of the two reference genes was stable according to BestKeeper software (Pfaffl et al., 2004). *Actin* was not used as an endogenous control gene in the experiments with adult bees because its expression was affected by the treatments.

To investigate whether synthetic queen mandibular pheromone (QMP) affects *lncov1* and/or *Tudor-SN* expression, newly-emerged workers from four source colonies headed by naturally-mated queens were housed in cages (n = 8 cages, 4 QMP^+^ and 4 QMP^-^,150 bees per cage) for four days at 34 °C [see (Cardoso-Júnior et al., 2020) for further details]. QMP^+^ cages contained a 0.5 queen equivalent released per day from a QMP strip (Phero Tech Inc., Canada), which is an effective queen mimic in cage experiments with young workers (Cardoso-Júnior et al., 2020, 2021a). QMP^-^ cages contained no QMP strip. Pollen, honey and water were provided *ad libitum*. Food was replenished when necessary. Workers collected on Day 0 (directly from the brood comb), Day 1, and Day 4 were immediately put on dry ice to determine the basal expression (Day 0), the short-term response (Day 1), and the more long-term response (Day 4) to the QMP treatment. The ovaries were then dissected and gene expression was determined as described above with eight biological replicates sampled per colony, time point, and treatment (Table S6). One sample from colony B2QR Day 4 (Table S6) was removed from the RT-qPCR analyses because of its basal expression levels of both *lncov1* and *Tudor-SN*.

To quantify the expression of *lncov1* and *Tudor-SN* in queenright and queenless field colonies, we used cDNA libraries prepared for our prior study (Cardoso-Júnior et al., 2021b). Briefly, three host colonies and located at a remote Apiary at Crommelin Research station 100 km north of Sydney were split into queenright (QR) and queenless (QL) units. On the same day, brood frames from the three source colonies were placed in an incubator to obtain newly-emerged workers that were paint marked according to source colony. Source colonies were headed by single-drone inseminated queens, as previously described in (Cardoso-Júnior et al., 2021b). The QR and QL units were transferred to the apiary of University of Sydney and the newly-emerged, paint-marked workers (n = 400 per colony, 200 QR and 200 QL) were added to their respective colony pairs. After four days, marked workers were collected and stored at -80 °C. Ovaries were dissected, pooled (four pairs of ovaries per sample and Table S6) and total RNA was extracted. Gene expression was determined as above, with eight biological replicates sampled from each QR and QL colony pair (Table S6).

In addition, we tested whether a diet that promotes ovary activation (royal jelly) affects the expression of *lncov1* and *Tudor-SN* in the ovaries of caged workers exposed or not to QMP. To do so, we used cDNA libraries from our previous study (Cardoso-Júnior et al., 2021a). Briefly, newly-emerged workers from two source colonies were randomly allocated to eight cages (4 QMP^+^ and 4 QMP^-^ each with 150 workers). In four of the cages the workers (2 QMP^+^ and 2 QMP^-^) received a diet composed of 50% honey and 50% royal jelly (Royal Jelly Australia, stored frozen), while those in the other four cages received a honey diet. Food was replenished when necessary. The cages were kept in an incubator at 34 °C in the dark for 7 days, when the bees were collected and snap frozen on dry ice and stored at -80 °C. Ovaries were dissected, and an ovary activation score based on a three-point scale (Ronai et al., 2016b). Results on the ovary activation scores are published elsewhere (Cardoso-Júnior et al., 2021a). Eight individual samples, each being a pool of four ovary pairs, were collected from each cage (Table S6), except for the activated ovaries of the royal jelly diet groups, which each sample represents a single pair of activated ovaries. With the ovary activation scores and gene expression data we performed correlation analyses (Fig. S3). RT-qPCR assays for the target genes of this experiment were done as described above.

### Sequence determination of *lncov1*

As the previously published *lncov1* sequence (Humann et al., 2013) had been generated by the assembly of 5’3’RACE amplicons from an uncharacterized EST (Humann and Hartfelder, 2011) it could not be deposited in full length in GenBank. Furthermore, the first published contig (Humann et al., 2013) contained a partial repeat within the *lncov1* sequence that was not present in the then available honey bee genome assembly. To obtain the full length of *lncov1* sequence and resolve the presence or not of the repeat sequence within *lncov1*, we obtained a full length *lncov1* product by amplification with primers containing a T7 promoter sequence at the 5’end of the reverse primer (Table S7) and Platinum *Taq* DNA Polymerase High Fidelity (Thermo Fisher Scientific, Waltham, MA) using the following conditions: 94 °C for 5 min, followed by 40 cycles of 94 °C for 30 s, 56 °C for 30 s, and 72 °C for 100 s, and a final elongation step at 72 °C for 7 min.

Amplicons were visualized by agarose gel electrophoresis (1%), purified with the Illustra GFX PCR DNA and Gel Band Purification Kit (GE Healthcare) and then cloned into the pGEM-T Easy Vector (Promega) for transformation of chemically-competent *E. coli* DH5α cells. We confirmed the completeness of the cloned *lncov1* sequence by Sanger sequencing of three independent clones. The final *lncov1* contig was assembled in CAP3 (Huang and Madan, 1999) software from the three forward and three reverse reads) and submitted to GenBank under the accession number OL505555.

*lncov1* has a 97% identity match, with only a 2 bp difference to its genomic scaffold position 11,932,549 – 11,933,630 (1082 bp) of linkage group 11 in the most recent *A. mellifera* genome assembly (Amel_HAv3.1) (Wallberg et al., 2019). We found no evidence of a repeat at its 5’end as originally reported (Humann et al., 2013). The full length *lncov1* sequence is now deposited in GenBank (accession number OL505555). We also confirmed the intronic location of *lncov1* in its host gene *LOC726407*, but due to changes in gene predictions in the most recent, chromosome-level genome assembly of *A. mellifera* (Wallberg et al., 2019), this is now the fifth intron.

### Whole-mount fluorescence *in situ* hybridisation (FISH)

To generate the *lncov1* riboprobes we used primers containing a T7 promoter sequence (Table S7). Amplification settings were 94 °C for 2 min, 40 cycles of 94 °C for 45 s, 60 °C for 45 s, 72 °C for 1 min, and a final extension step of 72 °C for 10 min. Amplicons were separated by agarose gel electrophoresis (1%), purified with the Illustra GFX PCR DNA and Gel Band Purification Kit (GE Healthcare), and then quantified spectrophotometrically. Subsequently, *lncov1* antisense and sense riboprobes (376 bp) labeled with Alexa Fluor™ 488 were produced by *in vitro* reverse transcription from the T7 promoter using a FISH Tag RNA Kit (Thermo Fisher Scientific, Waltham, MA).

Honey bee ovaries from L4, L5F3, and L5S3 worker larvae were processed as described for *Drosophila melanogaster* ovaries (Saunders and Cohen, 1999). After fixation in buffered heptane/paraformaldehyde [(82% heptane, 13.12%, HEPES buffer (0.1 M HEPES, pH 6.9; 2 mM MgSO_4_; and 1 mM EGTA), 0.66% paraformaldehyde and 1.64% DMSO)], the ovaries were washed with methanol and twice with ethanol 100%, and then stored in ethanol at -20 °C. Fixed ovaries were rehydrated in methanol, followed by methanol/PTw (PBS 1% and Tween 0.1%), and finally three times in PTw, and then transferred to a DMSO 1:9 PPTwT solution (paraformaldehyde 4%, PTw, and Triton X-100 0.1%) for 20 min at room temperature. After five washes in PTw, the ovaries were incubated for 30 s with protease K (40 mg/mL) and again washed with glycine (10 mg/mL) in PTw. Following two washes with PTw the ovaries were re-fixed with PPTwT and washed five times in PTw. Before hybridization, the ovaries were incubated for 10 min in PTw/Hybridization solution (HS: 50% formamide, 4x SSC, 50 mg/mL heparin, 1x Denhardt’s solution, 250 mg/mL yeast RNA and 500 mg/mL salmon testes DNA), and another 10 min with HS only. After 1 h at 45 °C in fresh HS, the fluorescent RNA riboprobe was added, and hybridzation proceeded for 16 h at 45 °C in the dark. The labeled ovaries were washed twice with HS and then, sequentially, with HS/PTw (3:1), HS/PTw (1:1), HS/PTw (1:3) and PTw. Nuclei were stained with DAPI/PTw (4000:1) before laser confocal in a TCS-SP5 System (Leica) microscopy (excitation laser set at 488 nm and emission at 525 nm), capturing optical sections of 0.5 or 1 µm thickness. Leica LAS-AF software was used for image acquisition and processing. No adjustments were made with respect to image brightness and/or contrast.

### Pull-down assays and mass spectrometry analysis

Recombinant pGEM-T easy vectors used to sequence the *lncov1* full-length were digested with *Eco*RI (Fermentas) and purified again with the Illustra GFX PCR DNA and Gel Band Purification Kit (GE Healthcare). *In vitro* reverse transcription was performed using the RiboMax T7 system (Promega, Madison, WI), and the product was biotinylated using the Pierce RNA 3’ End Desthiobiotinylation Kit (Thermo Fisher Scientific, Waltham, MA). Biotinylated *lncov1* fragments were linked to spherical beads of the Pierce Magnetic RNA-Protein Pull-Down Kit (Thermo Scientific), and the pull-down was performed according to the manufacturer’s protocol using whole-body protein extracts of L5F3 larvae. Proteins of these larvae were extracted using RIPA lysis buffer (0.75 M NaCl, 0,5% SDS, 0.25 M Tris, 5% Triton X-100, 100 mM EDTA supplemented with orthovanadate, 100 mM acid sodium pyrophosphate, 100 mM PMSF, 1% leupeptin), followed by centrifugation at 10,000 × *g* for 30 min at room temperature. Three independent pull-down experiments were performed, and the larval proteins bound to the *lncov1* pull-down beads were eluted following the manufacturer’s assay protocol.

The eluted proteins were analyzed by mass spectrometry using a shotgun proteomics approach (liquid chromatography-electrospray ionization-mass spectrometry, LC-ESI-MS/MS). After reduction with DTT (45 mM for 1 h at 56 °C), the proteins were alkylated with iodoacetamide (100 mM for 1 h at room temperature in a dark chamber) and digested with trypsin (Promega) diluted in ammonium bicarbonate (0.1 M for 24 h at 37 °C). Trypsination was stopped by the addition of formic acid, and the samples were stored at -20°C. For the mass spectrometry analysis, the samples were desalted by passage through a microcolumn containing reverse-phase resin (POROS R2, Perseptive Biosystems, USA), eluted in 60% methanol containing 5% formic acid, and dried by vacuum centrifugation. After resuspension in matrix solution (5 mg/mL 4-hydroxy cinnamic acid in 50% acetonitrile and 0.1% trifluoracetic acid), the samples were applied to the TOF plate of a MALDI-TOF/TOF-MS system (Axima Performance, Kratos-Shimadzu, Manchester, UK).

The spectrometry results were analyzed in the formats mascot generic and micromass (PKL) for MALDI-TOF/TOF and electrospray, respectively. The MASCOT MS/MS Ion Search tool (http://www.matrixscience.com) was used, and searches were done against the database NCBInr and Metazoan taxonomy using the following parameters: Carbamidomethyl (C) as fixed modifications, Deamidated (NQ) and Oxidation (M) as variable modifications, peptide tolerance ± 1.2 Da, MS/MS tolerance ± 0.8 Da, and peptide charge 1+.

### RNAi-mediated knockdown of *lncov1* and *Tudor-SN*

We designed primers flanked by the T7 sequence for *lncov1* and *Tudor-SN* to amplify 342 bp and 378 bp fragments of *lncov1* (i.e., ds-*lncov1*-I and ds-*lncov1*-II, respectively) and a 323 bp fragment of *Tudor-SN* (i.e., ds-*Tudor-SN*) (Table S7). PCR amplifications were performed using the following conditions: 95 °C for 3 min, followed by 40 cycles of 95 °C for 15 s, 60 °C for 30 s, and 72 °C for 1 min, and a final extension step of 72 °C for 7 min.

The respective amplicons were separated by agarose gel electrophoresis (1.2 %), purified with Illustra^TM^ DNA and Gel Band Purification Kit (GE Healthcare), and then cloned into pGEM®-T easy vector (Promega). Recombinant plasmids were used to transform chemically-competent *E. coli* DH5α cells. Positive clones were extracted with the QIAprep Spin Miniprep Kit (Qiagen, Hilden, Germany). Double-stranded RNAs (dsRNAs) were *in vitro* synthesised using the RiboMax T7 System Kit (Promega). As a control we prepared *GFP* dsRNA from a pGreen Lantern plasmid (Thermo Fisher Scientific) using specific primers (Table S7).

L5F2 worker larvae were collected from brood frames and transferred to plastic cups containing with 250 μL of a diet suitable for rearing worker larvae (Kaftanoglu et al., 2011) (53% royal jelly, 6% fructose, 6% glycose, 1% yeast extract, and 34% water). To the diet (250 μL) of each larva we added either 1 μg of *lncov1* dsRNA, 1 μg of *Tudor-SN* dsRNA, 1 μg of *GFP* dsRNA (negative control), or 1 µ L of water (untreated control). The larvae were kept in an incubator at 34 °C with controlled humidity for 24 hours before dissection of the ovaries. Three independent replicates were prepared for each of the treatment groups (*ds-GFP*, *ds-Tudor-SN,* or *ds-lncov-1*) and six for the untreated control group (see Table S6 for sample sizes). Knockdown efficiency was assessed using RT-qPCR assays with the respective primers for the target transcripts (*lncov1* or *Tudor-SN*). The primers for the RT-qPCR assays were designed so as to avoid overlap with the regions used for the dsRNA (Table S7). RT-qPCR assays were performed as described below.

As we did not achieve a reduction in *lncov1* expression via *in vivo* application of *ds-lncov1*, we next conducted an *in vitro* experiment. We prepared three independent pools of dissected ovaries from L5F2 worker larvae (n = 3), each consisting of 10 pairs of ovaries from L5F2 worker larvae, and cultivated these in Grace’s insect culture medium for 24 h at 34 °C. Ovaries cultured *in vitro* should require less dsRNA to achieve a successful knockdown, so we added *lncov1* dsRNA in the range from 10-1000 ng to the culture medium (200 μL per well). RT-qPCR assays for the target genes of this experiment were performed as described below.

### Effector caspase activity assay

Effector caspase activity was assessed using the Caspase-Glo 3/7 Assay System (Promega) following a previous protocol (Ronai et al., 2016b). The assay was performed on protein extracts from three independent pools of larval L5F3 ovaries of the *Tudor-SN* knockdown experiment, each pool consisting of ten pairs of ovaries. The luminescence signal representing caspase activity was detected in a SpectraMax L Microplate Luminometer (Promega). The luminescence was then normalized to the protein concentration of the sample, which was determined by Bradford assay.

### Juvenile hormone treatment of honey bee larvae

Fourth-instar worker larvae were topically treated with a 10 μg dose of Juvenile hormone III (Fluka, Buchs, Switzerland) dissolved in acetone (10 μg/μL), as previously described (23). Control larvae received an application of acetone (1 μL; solvent control), or were left untreated. Larvae were collected 6 h later, after they had molted to the L5F1 stage. Sample sizes of each group are listed in Table S6. The efficiency of the JH treatment was assessed by analysis of the honey bee *Krüppel homolog-1* gene, which is the immediate response gene for JH (Bellés, 2020). RT-qPCR assays for the target genes of this experiment were done as described below.

### *In silico* functional analyses of *lncov1* and Tudor-SN protein

The subcellular locations of *lncov1* and Tudor-SN protein were predicted using iLoc-LncRNA (http://lin-group.cn/server/iLoc-LncRNA/predictor.php) (Su et al., 2018) and ProtComp v9.0 (Softberry Inc., www.softberry.com/berry.phtml?topic=protcompan&group=programs&subgroup=proloc), respectively. Conserved domains encoded by *LOC726407* were identified by the CDART software (Geer et al., 2002) and Prosite Expasy (https://prosite.expasy.org/).

To investigate the expression of *lncov*1, *LOC726407*, *Tudor-SN* and *Gapdh* in published RNA-seq libraries of honey bee ovaries (Chen and Shi, 2020; Chen et al., 2017; Duncan et al., 2020), raw sequencing data was downloaded from Gene Expression Omnibus under the following accession codes: GSE120561, GSE93028, GSE119256. Raw reads were checked for quality with FastQC software v. 0.11.9 and when necessary, trimming was performed with Fastp v. 0.12.4. To quantify transcript abundance, a custom pipeline was developed (script is described in Supplementary Material 1). Briefly, we first indexed *lncov1* transcript to the list of transcripts known to be encoded by the *A. mellifera* genome (Amel_HAv3.1). The read count for each transcript was estimated probabilistically by Kallisto software (Bray et al., 2016) using a bootstrap of 1000x. This software repeats the transcript counting process in each library by resampling the data to increase accuracy in counting. This way, each bootstrap is considered a technical replication of a given biological sample. Normalized differential gene expression analyses across different libraries were performed with Sleuth software (Pimentel et al., 2017) in R (Team, 2018) using the Kallisto files as input. The pipeline developed for the Sleuth differential expression analysis is summarized in Supplementary Material 2. An adjusted q-value < 0.05 and a log_2_-fold change cutoff > 0.5, were considered for differential expression analyses. Gene expression of RNA-seq libraries between Chen et al. studies (Chen and Shi, 2020; Chen et al., 2017) were directly compared in this study due to their similarities regarding sample collection, RNA-seq library preparation and sequencing procedures.

### Evolutionary conservation of *lncov1*

The *lncov1* genomic sequence (1080 bp) was used as input for blastn searches against 38 hymenopteran genomes deposited in the Hymenoptera Genome Database – HGD (Elsik et al., 2018) (https://hymenoptera.elsiklab.missouri.edu/).

### Statistical analyses

For the experiments performed on larvae we used Kruskal-Wallis tests followed by Dunn’s post hoc test as the data distribution failed the Kolmogorov-Smirnov normality test. The only exception was with the JH-III treatment (Fig. S2D-F), where One-Way ANOVA with Bonferroni correction was performed, as it passed the Kolmogorov-Smirnov normality test. In the queen and worker comparison for *Tudor-SN* expression (Fig. 1I), Two-Way ANOVA with Bonferroni correction was applied, as the data followed a normal distribution (Kolmogorov-Smirnov normality test). For the experiments with adult honey bee workers, gene expression was analyzed as a dependent variable in a Generalized Linear Mixed Models (GLMM), with ‘colony’ as random effect, and ‘diet’, ‘QMP’, or ‘social context’ (queenright/queenless) as fixed effects. Two-tailed Student’s *t*-tests were used to analyze gene expression levels of *lncov1* and *Tudor-SN* in individual colonies (Figure 3A-B), or in activated and non-activated ovaries (insets of Figure 3C) as these data passed in the Kolmogorov-Smirnov normality test. For all analyses, a *p*-value < 0.05 was considered significant. Analyses were performed in GraphPad Prism 7, or in *R* (Team, 2018) using the packages lme4 (Bates et al., 2015) and lsmeans (Lenth, 2016).

### Material availability

All data supporting our findings are available in supplementary data files for this manuscript. *lncov1* full sequence is available in GenBank (accession number: OL505555).

### Data and Code availability

All material and data generated in this study can be directly retrieved from Lead Contact. Original codes used for transcriptome analyses of published RNA-seq datasets are provided in Material Supplementary 1 and 2.

## SUPPLEMENTAL INFORMATION

**Table S1.** Mascot hits for proteins identified by pulldown assays as binding partners to *Apis mellifera lncov1* RNA.

**Table S2.** Statistical details of caste-specific relative expression of *Tudor-SN*.

**Table S3.** Statistical details for the *Tudor-SN* knockdown *in vivo* experiment.

**Table S4.** Statistical details for the JH-III treatment *in vivo* experiment.

**Table S5.** Statistical details for experiments with adult bees.

**Table S6.** Detailed information of the biological samples collected and analyzed in this study.

**Table S7.** Primer list and their respective sequences, amplification temperature and reference.

**Table S8.** Results of BLASTn searches for *Apis mellifera lncov1* sequence as query in public Hymenoptera genomes databases.

**Supplementary Material 1.** Original scripts used to estimate transcript levels in RNA-seq libraries.

**Supplementary Material 2.** Original scripts used to perform differential gene expression in RNA-seq libraries.

