## Supplementary figures and images for "A long non-coding RNA is a key factor in the evolution of insect eusociality"

### Supplemental Figure 1

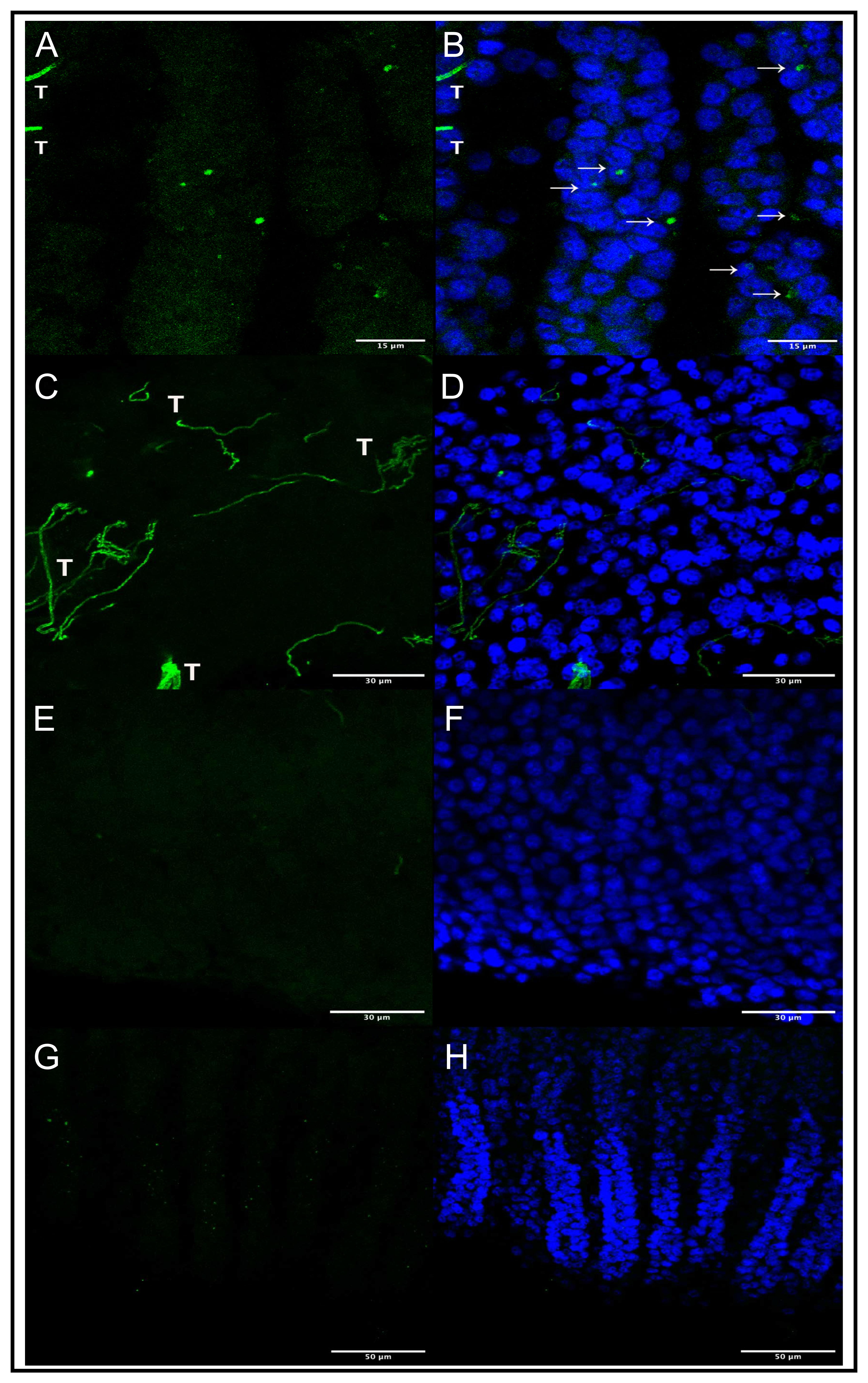

### Supplemental Figure 2

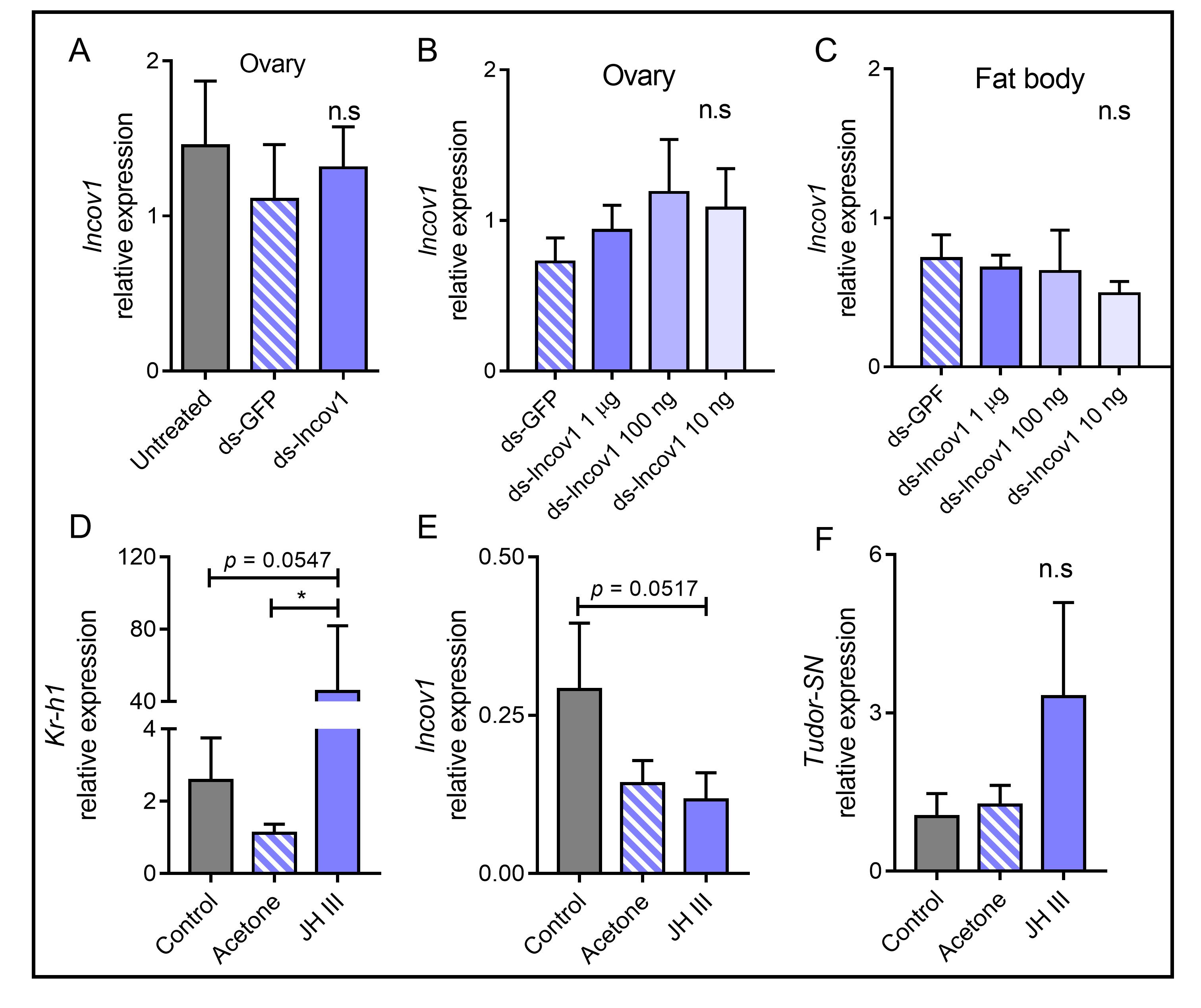

### Supplemental Figure 3

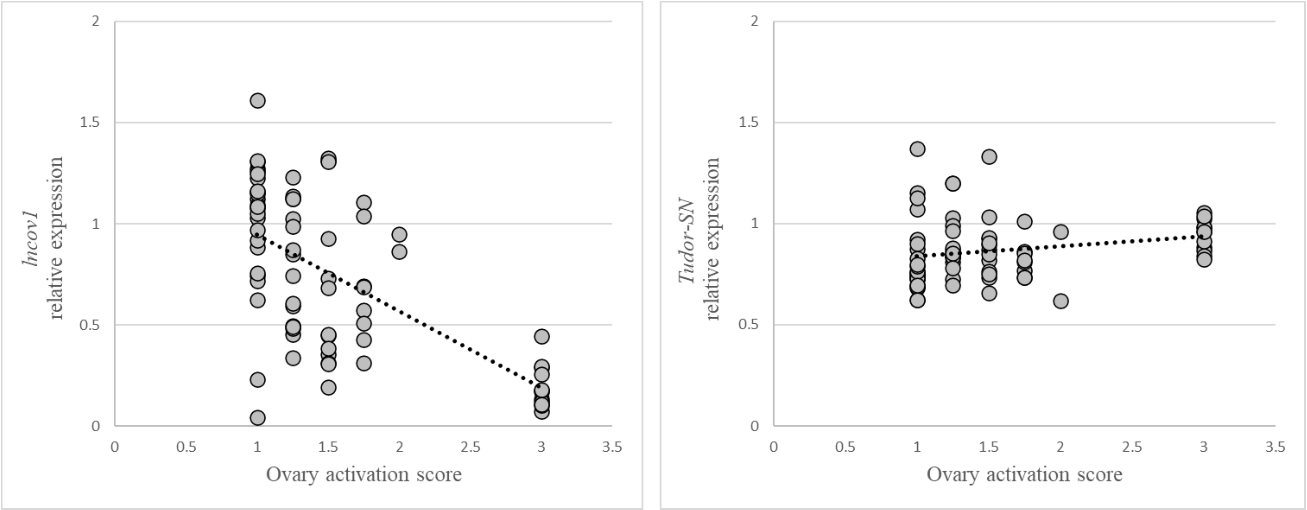

### Supplemental Figure 4

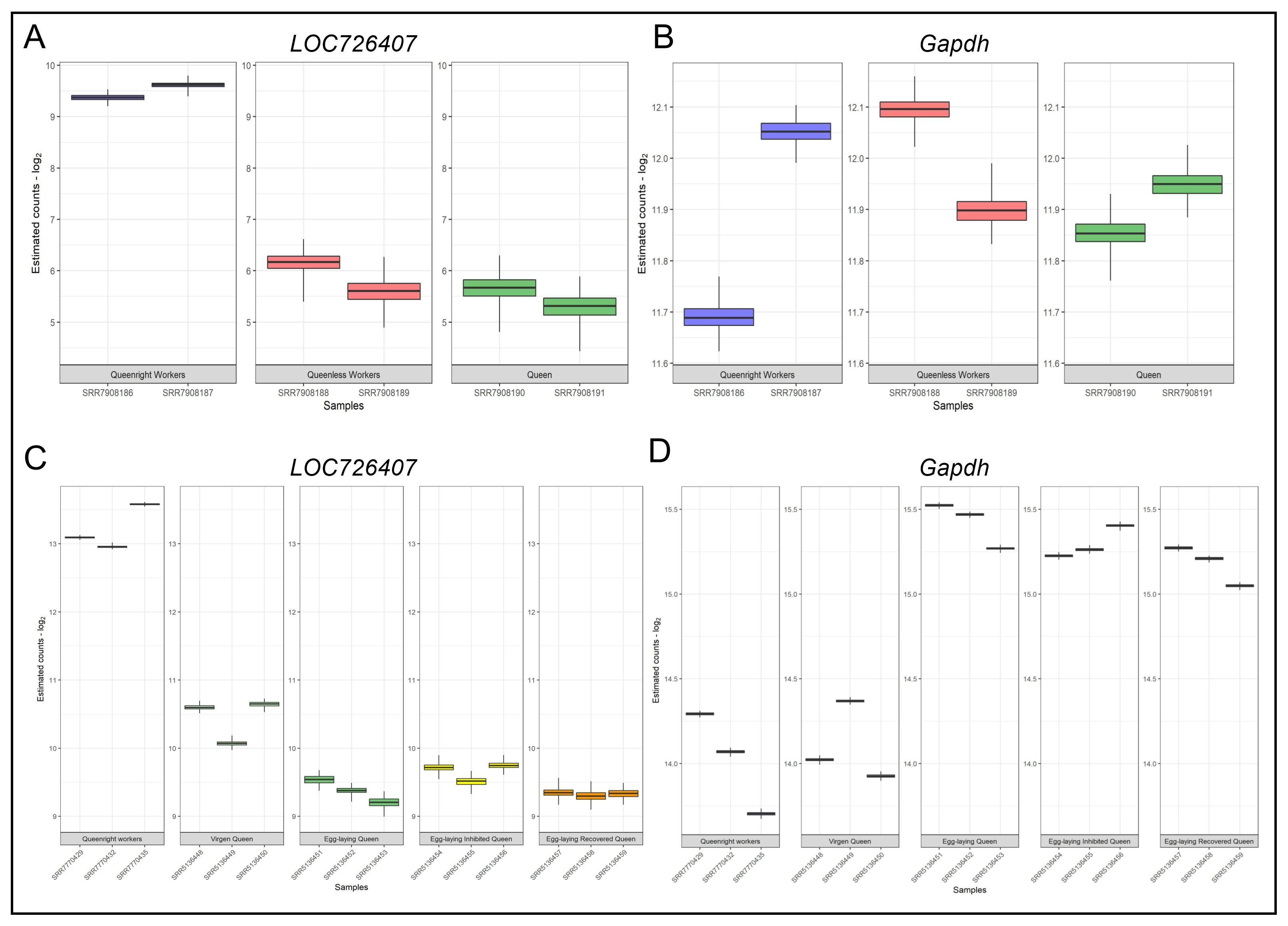

### Supplemental Figure 5

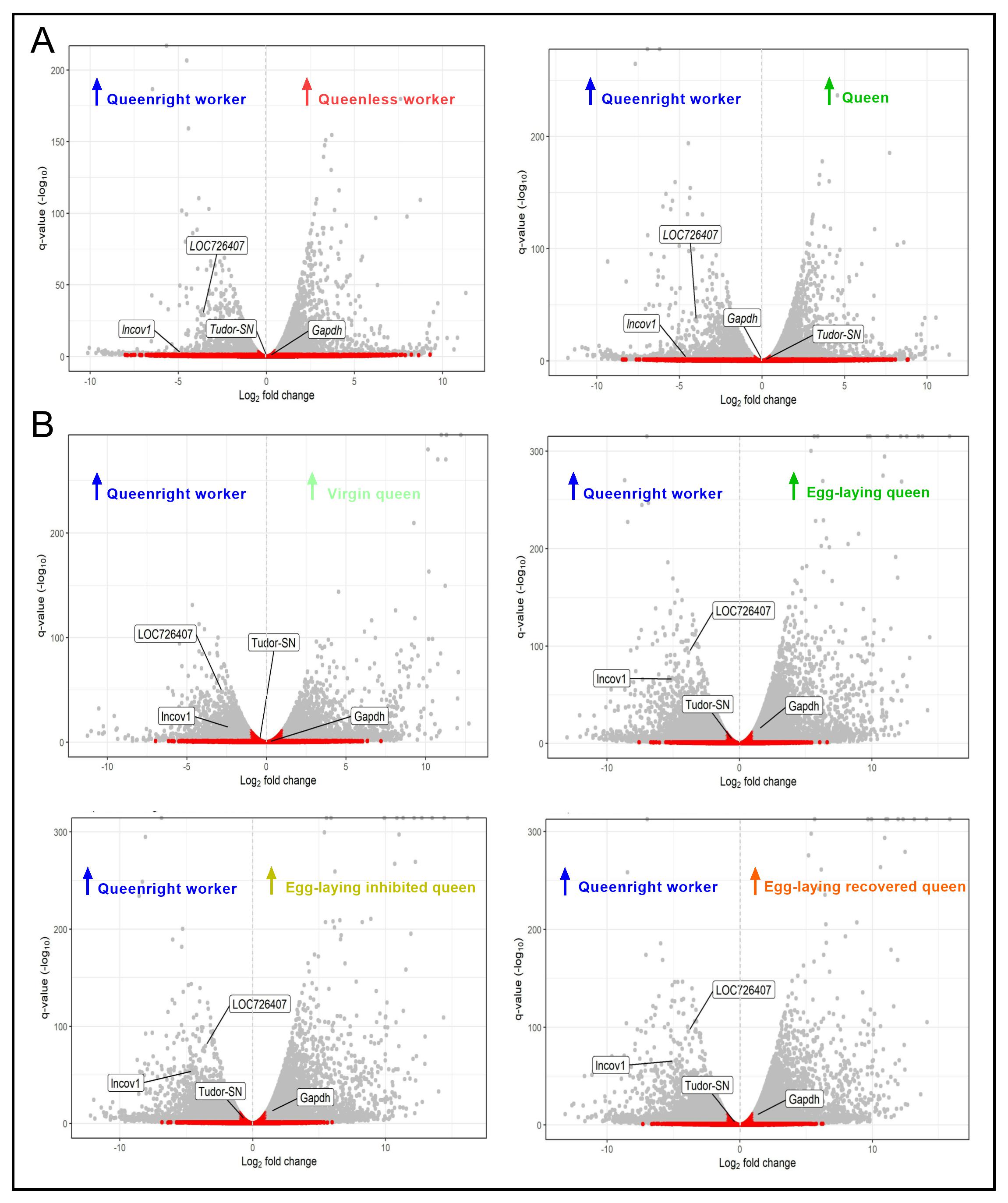
