## Supplemental Table 1 for "A long non-coding RNA is a key factor in the evolution of insect eusociality"

| **Table S1.** Mascot hits for proteins identified by pulldown assays as binding partners to *Apis mellifera lncov1* RNA. Shown are the NCBI and BeeBase protein IDs, the protein name, molecular mass, and the respective peptides identified by mass spectrometry. | | | |
| --- | --- | --- | --- |
| **Protein ID** | **Protein name** | **Molecular mass** | **Identified peptide** |
| gi\|328785330, XP_624638, GB40977 | Staphylococcal nuclease domain-containing protein 1 (*Apis mellifera*) | 101441 | LIGQDVAFVTEK |
|  |  |  | SSHYNLLQVAESK |
|  |  |  | GLISDGLLLVQNQR |
|  |  |  | AVVEFVTSGSR |
|  |  |  | LIGQDVAFVTEK |
|  |  |  | GNIAEILLSEGFAK |
|  |  |  | SSHYNLLQVAESK |
|  |  |  | GLISDGLLLVQNQR |
|  |  |  | LNNNVTAVTLVDSSSNEDIAK |
| gi\|66565249, XP_62027, GB47103 | Elongation factor 1-beta~ (*Apis mellifera*) | 24657 | TPVILGNNIAAGK |
|  |  |  | TPVILGNNIAAGK |
|  |  |  | SSVVLDVKPWDDETDMK |
| gi\|571561871, XP_006566786, GB45286 | Elongation factor 1-gamma-like (*Apis mellifera*) | 49240 | GNFDLDDFKR |
|  |  |  | ALTALNSHLLTR |
|  |  |  | VLIAAQYSGAQIK |
| gi\|571549351, XP_006561841, GB42862 | Elongation factor 1-delta isoform X2 (*Apis mellifera*) | 29935 | DILDSSVFQELK |
|  |  |  | KIETDGLLWGASK |
|  |  |  | SNIILDVKPWDDETDMK |
|  |  |  | GNQQTTSPILSAGGSLANEVAK |
| gi\|769840105, XP_011631225 | Elongation factor 1-delta isoform X1 (*Pogonomyrmex barbatus*) | 50929 | KIETDGLIWGASK |
|  |  |  | SNIILDVKPWDDETDMK |
| gi\|48097100, XP_391843, GB50979 | 3-ketoacyl-CoA thiolase, mitochondrial-like isoform X2 (*Apis mellifera*) | 42706 | NATDLSVIAAK |
|  |  |  | LNVDGGSIALGHPLAASGSR |
|  |  |  | LNVDGGSIALGHPLAASGSR |
|  |  |  | VDNVIFGHVLPISSSDGGFLTR |
|  |  |  | VDNVIFGHVLPISSSDGGFLTR |
| gi\|297591983, NP_-1172073, GB50912 | 60S acidic ribosomal protein P1 (*Apis mellifera*) | 11673 | ELITNIGSGVGK |
