## Supplemental Table 2 for "A long non-coding RNA is a key factor in the evolution of insect eusociality"

| **Table S2.** Statistical details of caste-specific relative expression of *Tudor-SN*. Significant results (*p* <0.05) are bolded. | | |
| --- | --- | --- |
| **Test** | **Group** | ***Tudor-SN*** |
| Two-Way ANOVA | Age | **F_(4,18)_ = 26.49 *p* < 0.0001** |
|  | Caste | **F_(1,18)_ = 6.166 *p* = 0.023** |
|  | Interaction: Age*Caste | **F_(4,18)_ = 5.357 *p* = 0.005** |
| Least square comparison after Bonferroni correction | L4 (Queen vs. Worker) | *t* = 1.7, DF = 18, *p* > 0.05 |
|  | L5F1 (Queen vs. Worker) | *t* = 2.01, DF = 18, *p* > 0.05 |
|  | L5F2 (Queen vs. Worker) | *t* = 0.64, DF = 18, *p* > 0.05 |
|  | L5F3 (Queen vs. Worker) | ***t* = 4.34, DF = 18, *p <* 0.01** |
|  | L5S3 (Queen vs. Worker) | *t* = 0.06, DF = 18 *p* > 0.05 |
