## Supplemental Table 3 for "A long non-coding RNA is a key factor in the evolution of insect eusociality"

| **Table S3.** Statistical details for the *Tudor-SN* knockdown *in vivo* experiment. Significant results are bolded. | | | |
| --- | --- | --- | --- |
| Target gene | Kruskal-Wallis test | Dunn’s post hoc test | |
| *ds-Tudor-SN* |  | *ds-Tudor-SN vs.* Control | *ds-Tudor-SN* *vs. dsGFP* |
| *Tudor-SN* | **H_(2)_ = 6.269 *p* = 0.028** | ***p* < 0.05** | *p* > 0.05 |
| *lncov1* | **H_(2)_ = 7.467 *p* = 0.07** | ***p* < 0.05** | *p* > 0.05 |
| *Anarchy* | H_(2)_ = 0.628 *p* = 0.77 | *p* > 0.05 | *p* > 0.05 |
| *Ark* | **H_(2)_ = 5.654 *p* = 0.046** | ***p* < 0.05** | *p* > 0.05 |
| *Buffy* | **H_(2)_ = 6.231 *p* = 0.033** | *p* > 0.05 | *p* > 0.05 |
| *Caspase-3* | H_(2)_ = 1.962 *p* = 0.41 | *p* > 0.05 | *p* > 0.05 |
| Caspase-3 activity | **H_(2)_ = 5.956 *p* = 0.025** | ***p* < 0.05** | *p* > 0.05 |
| *ds-lncov1* | Kruskal-Wallis Test | *ds-lncov1 vs.* Control | *ds-lncov1* *vs. dsGFP* |
| *lncov1 (ovary)* | H_(2)_ = 1.689 *p* = 0.51 | *p* > 0.05 | *p* > 0.05 |
| *lncov1 (fat body)* | H_(2)_ = 1.462 p = 0.74 | *p* > 0.05 | *p* > 0.05 |
