## Supplemental Table 4 for "A long non-coding RNA is a key factor in the evolution of insect eusociality"

| **Table S4.** Statistical details for the JH-III treatment *in vivo* experiment. Significant results are bolded. | | | |
| --- | --- | --- | --- |
| Target gene | One-Way ANOVA test | Bonferroni’s post hoc test | |
|  |  | Acetone *vs.* JH-III | Control *vs.* JH-III |
| *Kr-h1* | **F_(2)_ = 7.75 *p* = 0.0207** | ***p =* 0.0146** | *p* = 0.0547 |
| *lncov1* | F_(2)_ = 2.795 *p* = 0.0803 | *p =* 0.436 | *p* = 0.0524 |
| *Tudor-SN* | F_(2)_ = 0.85 *p* = 0.44 | *p* = 0.636 | *p* = 0.48 |
