## Supplemental Table 5 for "A long non-coding RNA is a key factor in the evolution of insect eusociality"

| T**able S5.** Statistical details for experiments with adult bees. Shown are the results for the QMP experiment (Fig. 3A), the results for queen presence/absence experiment (Fig. 3B), and the results for the royal jelly feeding experiment (Fig 3C). Significant results (*p* < 0.05) are bolded. | | | |
| --- | --- | --- | --- |
| **QMP experiment** | **Group** | ***lncov1*** | ***Tudor-SN*** |
| Main effects: GLMM test | QMP | **X^2^ = 7.8, *p* = 0.005** | **X^2^ = 12.96, *p* = 3.1E-4** |
|  | Age | **X^2^ = 51.85, *p* = 5.9E-13** | **X^2^ = 53.17, *p* = 3E-13** |
|  | Interaction: QMP*Age | X^2^ = 1.92, *p* = 0.16 | X^2^ = 2.78, *p* = 0.095 |
| Least square comparison after Bonferroni correction | Day 1 (QMP^+^ vs. QMP^-^) | *p* = 0.39 | *p* = 0.21 |
|  | Day 4 (QMP^+^ vs. QMP^-^) | ***p* = 0.006** | ***p* = 5E-4** |
| **Queen presence experiment** | **Group** | ***lncov1*** | ***Tudor-SN*** |
| Main Effect: GLMM test | Queen (Queenright vs. Queenless) | **X^2^ = 17.71, *p* = 2.5E-5** | **X^2^ = 16.27, *p* = 5.4E-5** |
| Two-tailed Student's *t*-test | Colony A (Queenright vs. Queenless) | ***t_(14_*_)_ = 2.62, *p* = 0.02** | ***t_(14_*_)_ = 6.7, *p* <0.0001** |
|  | Colony B (Queenright vs. Queenless) | ***t_(14_*_)_ = 2.89, *p* = 0.011** | *t_(14_*_)_ = 1.73, *p* = 0.105 |
|  | Colony C (Queenright vs. Queenless) | ***t_(14_*_)_ = 4.71, *p* = 3E-4** | *t_(14_*_)_ = 1.17, *p* = 0.25 |
| **Royal jelly experiment** | **Group** | ***lncov1*** | ***Tudor-SN*** |
| Main effects: GLMM test | QMP | **X^2^ = 7.61, *p* = *0.005*** | **X^2^ = 25.2, *p* = 5.16E-7** |
|  | Diet | **X^2^ = 12.36, *p* = 4.3E-4** | **X^2^ = 5.02, *p* = 0.024** |
|  | Interaction: QMP*Diet | X^2^ = 0.0002, *p* = 0.98 | **X^2^ = 14.21, *p* = 1.62E-4** |
| Least square comparison after Bonferroni correction | QMP^-^/Control vs. QMP^+^/Control | ***p* = 0.036** | ***p* < 0.0001** |
|  | QMP^-^/Control vs. QMP^-^/Royal Jelly | ***p* = 0.004** | *p* = 0.12 |
|  | QMP^-^/Control vs. QMP^+^/Royal Jelly | *p* = 0.97 | *p* = 0.16 |
|  | QMP^+^/Control vs. QMP^-^/Royal Jelly | ***p* < 0.0001** | ***p* = 0.007** |
|  | QMP^+^/Control vs. QMP^+^/Royal Jelly | ***p* = 0.009** | ***p* = 0.016** |
|  | QMP^-^/Royal Jelly vs. QMP^+^/Royal Jelly | ***p* = 0.012** | *p* = 1 |
| Two-tailed Student's *t*-test | Active ovaries (QMP^-^/Royal Jelly) vs. inactive ovaries (QMP^-^/Royal Jelly) | ***t_(_*_26)_ = 3.92, *p* = 6E-4** | ***t_(_*_26)_ = 2.88, *p* = 0.0078** |
