## Supplemental Table 6 for "A long non-coding RNA is a key factor in the evolution of insect eusociality"

|  | **Table S6.** Detailed information of the biological samples collected and analyzed in this study. | | | | | | | | |
| --- | --- | --- | --- | --- | --- | --- | --- | --- | --- |
| **Experiment** | | **Figure** | **Caste** |  | **Developmental stage or treatment group** | **Tissue** | **Sample size** | **# tissues per sample** | **Notes** |
| Tissue-specific Relative expression of *lncov1* and *LOC726407* | | Fig 1A | Worker |  | L4 | Ovary | 3 | 10 | These samples were collected and published in: Humann et al. PLoS One 8, e78915 (2013) |
|  |  | Fig 1A | Worker |  | L5F1 | Ovary | 3 | 10 |  |
|  |  | Fig 1A | Worker |  | L5F2 | Ovary | 3 | 10 |  |
|  |  | Fig 1A | Worker |  | L5F3 | Ovary | 3 | 10 |  |
|  |  | Fig 1A | Worker |  | L5S1 | Ovary | 3 | 10 |  |
|  |  | Fig 1A | Worker |  | L5S2 | Ovary | 3 | 10 |  |
|  |  | Fig 1A | Worker |  | L5S3 | Ovary | 3 | 10 |  |
|  |  | Fig 1A | Worker |  | L4 | Fat body | 3 | 10 | RNA extraction for gene expression analyses |
|  |  | Fig 1A | Worker |  | L5F1 | Fat body | 3 | 10 |  |
|  |  | Fig 1A | Worker |  | L5F2 | Fat body | 3 | 10 |  |
|  |  | Fig 1A | Worker |  | L5F3 | Fat body | 3 | 10 |  |
|  |  | Fig 1A | Worker |  | L5S1 | Fat body | 3 | 10 |  |
|  |  | Fig 1A | Worker |  | L5S2 | Fat body | 3 | 10 |  |
|  |  | Fig 1A | Worker |  | L5S3 | Fat body | 3 | 10 |  |
|  |  | Fig 1A | Worker |  | L4 | Head | 3 | 10 |  |
|  |  | Fig 1A | Worker |  | L5F1 | Head | 3 | 10 |  |
|  |  | Fig 1A | Worker |  | L5F2 | Head | 3 | 10 |  |
|  |  | Fig 1A | Worker |  | L5F3 | Head | 3 | 10 |  |
|  |  | Fig 1A | Worker |  | L5S1 | Head | 3 | 10 |  |
|  |  | Fig 1A | Worker |  | L5S2 | Head | 3 | 10 |  |
|  |  | Fig 1A | Worker |  | L5S3 | Head | 3 | 10 |  |
|  |  | Fig 1A | Worker |  | L4 | Leg Imaginal discs | 3 | 4 |  |
|  |  | Fig 1A | Worker |  | L5F1 | Leg Imaginal discs | 3 | 4 |  |
|  |  | Fig 1A | Worker |  | L5F2 | Leg Imaginal discs | 3 | 4 |  |
|  |  | Fig 1A | Worker |  | L5F3 | Leg Imaginal discs | 3 | 4 |  |
|  |  | Fig 1A | Worker |  | L5S1 | Leg Imaginal discs | 3 | 4 |  |
|  |  | Fig 1A | Worker |  | L5S2 | Leg Imaginal discs | 3 | 4 |  |
|  |  | Fig 1A | Worker |  | L5S3 | Leg Imaginal discs | 3 | 4 |  |
| Caste-specific expression of *Tudor-SN* | | Fig 1I | Worker |  | L4 | Ovary | 3 | 10 | RNA extraction for gene expression analyses |
|  |  | Fig 1I | Worker |  | L5F1 | Ovary | 3 | 10 |  |
|  |  | Fig 1I | Worker |  | L5F2 | Ovary | 3 | 10 |  |
|  |  | Fig 1I | Worker |  | L5F3 | Ovary | 3 | 10 |  |
|  |  | Fig 1I | Worker |  | L5S3 | Ovary | 3 | 10 |  |
|  |  | Fig 1I | Queen |  | L4 | Ovary | 3 | 10 |  |
|  |  | Fig 1I | Queen |  | L5F1 | Ovary | 3 | 10 |  |
|  |  | Fig 1I | Queen |  | L5F2 | Ovary | 3 | 10 |  |
|  |  | Fig 1I | Queen |  | L5F3 | Ovary | 3 | 10 |  |
|  |  | Fig 1I | Queen |  | L5S3 | Ovary | 3 | 10 |  |
| RNAi *in vivo* | | Figs 2A-F | Worker |  | L5F3 - Untreated | Ovary | 3 | 10 | RNA extraction for gene expression analyses |
|  |  | Figs. 2A-F | Worker |  | L5F3 - ds-Tudor-SN - 1μg | Ovary | 3 | 10 |  |
|  |  | Figs. 2A-F | Worker |  | L5F3 - ds-GFP - 1μg | Ovary | 3 | 10 |  |
|  |  | Fig. S2A | Worker |  | L5F3 - Untreated | Ovary | 3 | 10 |  |
|  |  | Fig. S2A | Worker |  | L5F3 - ds-lncov1 - 1μg | Ovary | 3 | 10 |  |
|  |  | Fig. S2A | Worker |  | L5F3 - ds-GFP - 1μg | Ovary | 3 | 10 |  |
|  |  | Fig 2G | Worker |  | L5F3 - Untreated | Ovary | 3 | 10 | Protein extraction for Caspase-3 activity assay |
|  |  | Fig 2G | Worker |  | L5F3 - ds-Tudor-SN - 1μg | Ovary | 3 | 10 |  |
|  |  | Fig 2G | Worker |  | L5F3 - ds-GFP - 1μg | Ovary | 3 | 10 |  |
| RNAi *in vitro* | | Fig S2B | Worker |  | L5F3 - ds-lncov1 - 1μg | Ovary | 3 | 10 | RNA extraction for gene expression analyses |
|  |  | Fig S2B | Worker |  | L5F3 - ds-lncov1 - 100ng | Ovary | 3 | 10 |  |
|  |  | Fig S2B | Worker |  | L5F3 - ds-lncov1 - 10ng | Ovary | 3 | 10 |  |
|  |  | Fig S2B | Worker |  | L5F3 - ds-GFP - 1μg | Ovary | 3 | 10 |  |
|  |  | Fig S2C | Worker |  | L5F3 - ds-lncov1 - 1μg | Fat body | 3 | 10 |  |
|  |  | Fig S2C | Worker |  | L5F3 - ds-lncov1 - 100ng | Fat body | 3 | 10 |  |
|  |  | Fig S2C | Worker |  | L5F3 - ds-lncov1 - 10ng | Fat body | 3 | 10 |  |
|  |  | Fig S2C | Worker |  | L5F3 - ds-GFP - 1μg | Fat body | 3 | 10 |  |
| Juvenile hormone treatment | | Figs S2D-F | Worker |  | L5F2 - Control | Ovary | 9 | 10 | RNA extraction for gene expression analyses |
|  |  | Figs S2D-F | Worker |  | L5F2 - Acetone | Ovary | 9 | 10 |  |
|  |  | Figs S2D-F | Worker |  | L5F2 - HJ-III | Ovary | 6 | 10 |  |
| QMP treatment | | Fig 3A | Worker |  | Day 0 | Ovary | 32 | 4 | Four independent colonies were analyzed. For each colony, eight independent biological samples were collected, the exception being one sample from “B2QR” colony (Day 4), which was removed from RT-qPCR assays due to undetectable expression levels of *lncov1* and *Tudor-SN* |
|  |  | Fig 3A | Worker |  | Day 1 QMP^+^ | Ovary | 32 | 4 |  |
|  |  | Fig 3A | Worker |  | Day 1 QMP^-^ | Ovary | 32 | 4 |  |
|  |  | Fig 3A | Worker |  | Day 4 QMP^+^ | Ovary | 32 | 4 |  |
|  |  | Fig 3A | Worker |  | Day 4 QMP^-^ | Ovary | 31 | 4 |  |
| Queenright *vs.* Queenless - field experiment | | Fig 3B | Worker |  | 4-days-old adult bees - Queenright | Ovary | 24 | 4 | Three colonies were analyzed, each one with eight independent biological samples of inactive ovaries |
|  |  | Fig 3B | Worker |  | 4-days-old adult bees - Queenless | Ovary | 24 | 4 |  |
| Royal Jelly diet | | Fig 3C | Worker |  | QMP^+^/Control Diet | Ovary | 16 | 4 | Two colonies were analyzed, each one with eight independent biological samples of inactive ovaries |
|  |  | Fig 3C | Worker |  | QMP^-^/Control Diet | Ovary | 16 | 4 |  |
|  |  | Fig 3C | Worker |  | QMP^+^/Royal jelly Diet | Ovary | 18 | 4 | Two colonies were analyzed, each one with eight independent biological samples of inactive ovaries plus two samples of activated ovaries |
|  |  | Fig 3C | Worker |  | QMP^-^/Royal jelly Diet | Ovary | 26 | 4 | Two colonies were analyzed, the first with nine independent biological sample of inactive ovaries plus two samples of activated ovaries; the second colony with eight independent biological samples of inactive ovaries plus seven samples of activated ovaries |
