## Supplemental Table 6 for "A long non-coding RNA is a key factor in the evolution of insect eusociality"

**Table S7.** Primer list and their respective sequences, amplification temperature and reference. T7 sequence is underlined.

| **Experiment** | **Target** | **Primer** | **Primer sequence** | **Reference** |
| --- | --- | --- | --- | --- |
| Gene expression | *lncov1* (intron 6 of *LOC726407*) | F | GGAGAAGCTTTGGGGAGAG | 1 |
|  |  | R | CTGCTACACACCACCATAAC |  |
|  | *LOC726407* (GB45056) | F | CGAGGTAGCGGTTCCTACAAG | 1 |
|  |  | R | AATGCTTCTGTCCGGAACAT |  |
|  | *Tudor-SN* (GB40977) | F | GTTACCAGTGGATCGCGTCT | This study |
|  |  | R | AGCAATGCTCTCGGGTGAAA |  |
|  | *Ark* (GB52453) | F | GTTTGTGCCAGTATGACTGA | 2 |
|  |  | R | CCAATATGTGTCCAAAGAAGAA |  |
|  | *Buffy* (GB49154) | F | GGTATTGCCGTGGATTGTGT | 2 |
|  |  | R | CAGATCTGTATCGAGTTGCTA |  |
|  | *Caspase-3* (GB41369) | F | CATCACAAACGAAGAGGCAC | 3 |
|  |  | R | GTCGACGATGAGTTCCACG |  |
|  | *Anarchy* (GB48961) | F | ACAAAGGAATGGAAGCCAAA | 4 |
|  |  | R | GGAAATTTACACGGGGAACA |  |
|  | *Kruppel-homolog 1* (GB45427) | F | GCACTGGCAGTGACAAGGAA | 5 |
|  |  | R | CGTGGAGTGTTATCGTAAGTAGCAA |  |
|  | *Rpl32* (*Rp49;* GB47227) | F | CGTCATATGTTGCCAACTGGT | 6 |
|  |  | R | TTGAGCACGTTCAACAATGG |  |
|  | *Actin* (GB44311) | F | TGCCAACACTGTCCTTTCTG | 6 |
|  |  | R | AGAATTGACCCACCAATCCA |  |
|  | *Ef1α* (GB52028) | F | TGCAACCTACTAAGCCGATG | 4 |
|  |  | R | GACCTTGCCCTGGGTATCTT |  |
| Pulldown and Sanger sequencing | *lncov1* (intron 6 of *LOC726407*) | F | GAGAGGAGAAGCTTTGGG | This study |
|  |  | R-T7 | TAATACGACTCACTATAGGGCGACTTTATGAAAGAATATTGCATATTCCTC |  |
| RNAi | *lncov1* (intron 6 of *LOC726407*) II | F-T7 | TAATACGACTCACTATAGGGCGAactgagtttcgggtgtgagg | This study |
|  |  | R-T7 | TAATACGACTCACTATAGGGCGAcaataagcggggataaccaa |  |
|  | *lncov1* (intron 6 of *LOC726407*) I | F-T7 | TAATACGACTCACTATAGGGCGAtggttatccccgcttattgg | This study |
|  |  | R-T7 | TAATACGACTCACTATAGGGCGActttgatgcttgcgtttgacgg |  |
|  | *Tudor-SN* (GB40977) | F-T7 | TAATACGACTCACTATAGGGCGACATATTCTCGCTTGTGTTGCGC | This study |
|  |  | R-T7 | TAATACGACTCACTATAGGGCGATCGTCTGCTCGTATGTCACC |  |
|  | *GFP* (pGreen Lantern - Thermo Fisher Scientific) | F-T7 | TAATACGACTCACTATAGGGCGACACTGGAGTGGTCCCAGTTCT | This study |
|  |  | R-T7 | TAATACGACTCACTATAGGGCGATGGGTATCTTGAGAAGCATTGAA |  |
| FISH | *lncov1* antisense | F | CTGAGTTTCGGGTGTGAGG | 1 |
|  |  | R-T7 | TAATACGACTCACTATAGGGCAATAAGCGGGGATAACCAA |  |
|  | *lncov1* sense | F-T7 | TAATACGACTCACTATAGGGCTGAGTTTCGGGTGTGTGAGG | 1 |
|  |  | R | CAATAAGCGGGGATAACCAA |  |
