## Supplemental Material 1 for "A long non-coding RNA is a key factor in the evolution of insect eusociality": Supplementary material 1.html

 transcriptomeAnalysis\_kallisto-sleuth

### Transcriptome Analysis with kallisto

###### Autor: Luiz Carlos Vieira

###### Date: 25/03/2022

Kallisto rely on a pseudoalignment which does not identify the positions of the reads in the transcripts, only their potential
transcripts of origin. It avoids doing an alignment of each read to a reference genome, using just the transcriptome sequences
as reference genome [1].

Standard methods for quantifying expression relied on mapping sequenced reads to a reference genome, assigning reads to a position
in the genome and the gene or isoform expression values are generated by counting the number of overlapping reads (overlapping the
features of interest) [2].

#### Downloading Rna-Seq datasets

```
# GSE120561 sample experiment id
# Overall design: RNA-seq to measure gene expression in queen,
# queen right worker and actively laying worker ovaries
# tissue: ovary
# Library strategy:   RNA-Seq
# Library source: transcriptomic
# Library selection:  cDNA
# Instrument model:   Illumina HiSeq 2500

curl -L ftp://ftp.sra.ebi.ac.uk/vol1/fastq/SRR790/006/SRR7908186/SRR7908186.fastq.gz -o SRR7908186_W1_Worker_Pool_1.fastq.gz
curl -L ftp://ftp.sra.ebi.ac.uk/vol1/fastq/SRR790/007/SRR7908187/SRR7908187.fastq.gz -o SRR7908187_W2_Worker_Pool_2.fastq.gz
curl -L ftp://ftp.sra.ebi.ac.uk/vol1/fastq/SRR790/008/SRR7908188/SRR7908188.fastq.gz -o SRR7908188_AW1_Active_Pool_1.fastq.gz
curl -L ftp://ftp.sra.ebi.ac.uk/vol1/fastq/SRR790/009/SRR7908189/SRR7908189.fastq.gz -o SRR7908189_AW2_Active_Pool_2.fastq.gz
curl -L ftp://ftp.sra.ebi.ac.uk/vol1/fastq/SRR790/001/SRR7908191/SRR7908191.fastq.gz -o SRR7908191_Q2_Queen_Pool_2.fastq.gz
curl -L ftp://ftp.sra.ebi.ac.uk/vol1/fastq/SRR790/000/SRR7908190/SRR7908190.fastq.gz -o SRR7908190_Q1_Queen_Pool_1.fastq.gz

# ----------------------------------------------------------------------------------------------------------------------------

# GSE93028 sample experiment id
# Title:    Integration of lncRNA-miRNA-mRNA reveals novel insights into reproductive regulation in honey bees
# Organism: Apis mellifera
# Experiment type:  Expression profiling by high throughput sequencing Non-coding RNA profiling by high throughput sequencing
# link: https://www.ncbi.nlm.nih.gov/geo/query/acc.cgi?acc=GSE93028

# Ovaries samples used in this study:  

# (1) ovaries of virgin queens (n=3); Characteristics: tissue: ovaries, age: virgin queen. 
curl -L ftp://ftp.sra.ebi.ac.uk/vol1/fastq/SRR513/008/SRR5136448/SRR5136448_1.fastq.gz -o SRR5136448_V01_mRNA_RNA-Seq_1.fastq.gz
curl -L ftp://ftp.sra.ebi.ac.uk/vol1/fastq/SRR513/008/SRR5136448/SRR5136448_2.fastq.gz -o SRR5136448_V01_mRNA_RNA-Seq_2.fastq.gz
curl -L ftp://ftp.sra.ebi.ac.uk/vol1/fastq/SRR513/009/SRR5136449/SRR5136449_1.fastq.gz -o SRR5136449_V02_mRNA_RNA-Seq_1.fastq.gz
curl -L ftp://ftp.sra.ebi.ac.uk/vol1/fastq/SRR513/009/SRR5136449/SRR5136449_2.fastq.gz -o SRR5136449_V02_mRNA_RNA-Seq_2.fastq.gz
curl -L ftp://ftp.sra.ebi.ac.uk/vol1/fastq/SRR513/000/SRR5136450/SRR5136450_1.fastq.gz -o SRR5136450_V03_mRNA_RNA-Seq_1.fastq.gz
curl -L ftp://ftp.sra.ebi.ac.uk/vol1/fastq/SRR513/000/SRR5136450/SRR5136450_2.fastq.gz -o SRR5136450_V03_mRNA_RNA-Seq_2.fastq.gz

# (2) ovaries of egg-laying queens (n=3); Characteristics tissue: ovaries, age: normal egg-laying queen
curl -L ftp://ftp.sra.ebi.ac.uk/vol1/fastq/SRR513/001/SRR5136451/SRR5136451_1.fastq.gz -o SRR5136451_Q01_mRNA_RNA-Seq_1.fastq.gz
curl -L ftp://ftp.sra.ebi.ac.uk/vol1/fastq/SRR513/001/SRR5136451/SRR5136451_2.fastq.gz -o SRR5136451_Q01_mRNA_RNA-Seq_2.fastq.gz
curl -L ftp://ftp.sra.ebi.ac.uk/vol1/fastq/SRR513/002/SRR5136452/SRR5136452_1.fastq.gz -o SRR5136452_Q02_mRNA_RNA-Seq_1.fastq.gz
curl -L ftp://ftp.sra.ebi.ac.uk/vol1/fastq/SRR513/002/SRR5136452/SRR5136452_2.fastq.gz -o SRR5136452_Q02_mRNA_RNA-Seq_2.fastq.gz
curl -L ftp://ftp.sra.ebi.ac.uk/vol1/fastq/SRR513/003/SRR5136453/SRR5136453_1.fastq.gz -o SRR5136453_Q03_mRNA_RNA-Seq_1.fastq.gz
curl -L ftp://ftp.sra.ebi.ac.uk/vol1/fastq/SRR513/003/SRR5136453/SRR5136453_2.fastq.gz -o SRR5136453_Q03_mRNA_RNA-Seq_2.fastq.gz

# (3) ovaries of egg-laying inhibited queens (n=3); Characteristics tissue: ovaries, age: egg-laying inhibited queen treatment: Caged for 7 days
curl -L ftp://ftp.sra.ebi.ac.uk/vol1/fastq/SRR513/004/SRR5136454/SRR5136454_1.fastq.gz -o SRR5136454_C01_mRNA_RNA-Seq_1.fastq.gz
curl -L ftp://ftp.sra.ebi.ac.uk/vol1/fastq/SRR513/004/SRR5136454/SRR5136454_2.fastq.gz -o SRR5136454_C01_mRNA_RNA-Seq_2.fastq.gz
curl -L ftp://ftp.sra.ebi.ac.uk/vol1/fastq/SRR513/005/SRR5136455/SRR5136455_1.fastq.gz -o SRR5136455_C02_mRNA_RNA-Seq_1.fastq.gz
curl -L ftp://ftp.sra.ebi.ac.uk/vol1/fastq/SRR513/005/SRR5136455/SRR5136455_2.fastq.gz -o SRR5136455_C02_mRNA_RNA-Seq_2.fastq.gz
curl -L ftp://ftp.sra.ebi.ac.uk/vol1/fastq/SRR513/006/SRR5136456/SRR5136456_1.fastq.gz -o SRR5136456_C03_mRNA_RNA-Seq_1.fastq.gz
curl -L ftp://ftp.sra.ebi.ac.uk/vol1/fastq/SRR513/006/SRR5136456/SRR5136456_2.fastq.gz -o SRR5136456_C03_mRNA_RNA-Seq_2.fastq.gz

# (4) ovaries of egg-laying recovered queens (n=3); Characteristics tissue: ovaries, age: egg-laying recovered queen
# treatment: the queen was released after caged for seven days. And on the eighth day, the queen recovered to normal condition and laid eggs. 
curl -L ftp://ftp.sra.ebi.ac.uk/vol1/fastq/SRR513/007/SRR5136457/SRR5136457_1.fastq.gz -o SRR5136457_R01_mRNA_RNA-Seq_1.fastq.gz
curl -L ftp://ftp.sra.ebi.ac.uk/vol1/fastq/SRR513/007/SRR5136457/SRR5136457_2.fastq.gz -o SRR5136457_R01_mRNA_RNA-Seq_2.fastq.gz
curl -L ftp://ftp.sra.ebi.ac.uk/vol1/fastq/SRR513/008/SRR5136458/SRR5136458_1.fastq.gz -o SRR5136458_R02_mRNA_RNA-Seq_1.fastq.gz
curl -L ftp://ftp.sra.ebi.ac.uk/vol1/fastq/SRR513/008/SRR5136458/SRR5136458_2.fastq.gz -o SRR5136458_R02_mRNA_RNA-Seq_2.fastq.gz
curl -L ftp://ftp.sra.ebi.ac.uk/vol1/fastq/SRR513/009/SRR5136459/SRR5136459_1.fastq.gz -o SRR5136459_R03_mRNA_RNA-Seq_1.fastq.gz
curl -L ftp://ftp.sra.ebi.ac.uk/vol1/fastq/SRR513/009/SRR5136459/SRR5136459_2.fastq.gz -o SRR5136459_R03_mRNA_RNA-Seq_2.fastq.gz

# ----------------------------------------------------------------------------------------------------------------------------

# GSE119256 sample experiment id
# Title:    Genome-wide analysis of coding and non-coding RNAs in ovary of honey bee workers
# Organism: Apis mellifera
# Source name:  workers_ovary_mRNA including lncRNA
# Organism: Apis mellifera
# Characteristics social caste: honey bee workers
# tissue: ovaries
# molecule subtype: mRNA including lncRNA
# Extracted molecule    total RNA
# limk: https://www.ncbi.nlm.nih.gov/geo/query/acc.cgi?acc=GSE119256

curl -L ftp://ftp.sra.ebi.ac.uk/vol1/fastq/SRR777/009/SRR7770429/SRR7770429_1.fastq.gz -o SRR7770429_W01_mRNA_including_lncRNA_RNA-Seq_1.fastq.gz
curl -L ftp://ftp.sra.ebi.ac.uk/vol1/fastq/SRR777/009/SRR7770429/SRR7770429_2.fastq.gz -o SRR7770429_W01_mRNA_including_lncRNA_RNA-Seq_2.fastq.gz
curl -L ftp://ftp.sra.ebi.ac.uk/vol1/fastq/SRR777/002/SRR7770432/SRR7770432_1.fastq.gz -o SRR7770432_W02_mRNA_including_lncRNA_RNA-Seq_1.fastq.gz
curl -L ftp://ftp.sra.ebi.ac.uk/vol1/fastq/SRR777/002/SRR7770432/SRR7770432_2.fastq.gz -o SRR7770432_W02_mRNA_including_lncRNA_RNA-Seq_2.fastq.gz
curl -L ftp://ftp.sra.ebi.ac.uk/vol1/fastq/SRR777/005/SRR7770435/SRR7770435_1.fastq.gz -o SRR7770435_W03_mRNA_including_lncRNA_RNA-Seq_1.fastq.gz
curl -L ftp://ftp.sra.ebi.ac.uk/vol1/fastq/SRR777/005/SRR7770435/SRR7770435_2.fastq.gz -o SRR7770435_W03_mRNA_including_lncRNA_RNA-Seq_2.fastq.gz
```

#### Quality Control of reads

```
sample="GSE120561 GSE93028_GSE119256"; for i in $sample; do fastqc -t 4 ${i}/rawFastq/*.fastq -o /mnt/c/Users/luiz_/Downloads/transcriptomeAnalysis/${i}/QC; done

sample="GSE120561 GSE93028_GSE119256"; for i in $sample; do multiqc ${i}/QC/ -o ${i}/QC/
```

#### Adding lncov1 into the transcriptome fasta

```
cat ref_genome/Amel_HAv3.1_rna.fna kallisto/lncov1.fasta > kallisto/all_transcrits.fasta
```

#### Building up a index of transcripts FASTA sequences (kallisto index)

```
kallisto index -i kallisto/index_trans kallisto/all_transcrits.fasta
```

#### Verifying lncov1 among transcripts

```
makeblastdb -in kallisto/all_transcripts.fasta -dbtype nucl -input_type fasta -title "all_transcripts_Amel_HAv3.1" -out blast/db/all_transcripts_Amel_HAv3.1

blastn -db kallisto/all_transcripts_Amel_HAv3.1 -query kallisto/lncov1.fasta -out kallisto/res_blast_lncov1_x_all_trascripts.txt
```

Reference: Zheng Zhang, Scott Schwartz, Lukas Wagner, and Webb
Miller (2000), “A greedy algorithm for aligning DNA sequences”, J
Comput Biol 2000; 7(1-2):203-14.

Database: all\_transcripts\_Amel\_HAv3.1
27,888 sequences; 98,646,852 total letters

Query= lncov1 Clone\_01-F

Length=1152
Score E
Sequences producing significant alignments: (Bits) Value

lncov1 Clone\_01-F 2128 0.0

> lncov1 Clone\_01-F
> Length=1152

Score = 2128 bits (1152), Expect = 0.0
Identities = 1152/1152 (100%), Gaps = 0/1152 (0%)
Strand=Plus/Plus

#### Verifying strandness of reads

```
SAMPLE=GSE
KALLISTO_INDEX=/mnt/c/Users/luiz_/Downloads/transcriptomeAnalysis/index/index_trans
OUT_DIR=/mnt/c/Users/luiz_/Downloads/transcriptomeAnalysis/${SAMPLE}/kallisto
SEQ_DIR=/mnt/c/Users/luiz_/Downloads/transcriptomeAnalysis/${SAMPLE}/rawFastq

# Make a subset from a fastqc
zcat ${SEQ_DIR}/SRR7770429_W01_mRNA_including_lncRNA_RNA-Seq_1.fastq.gz | head -n16000 > ${SEQ_DIR}/test_1.fastq
zcat ${SEQ_DIR}/SRR7770429_W01_mRNA_including_lncRNA_RNA-Seq_2.fastq.gz | head -n16000 > ${SEQ_DIR}/test_2.fastq

# Running kallisto quant
kallisto quant -i ${KALLISTO_INDEX} -o ${OUT_DIR}/test_un ${SEQ_DIR}/test_1.fastq ${SEQ_DIR}/test_2.fastq
kallisto quant -i ${KALLISTO_INDEX} -o ${OUT_DIR}/test_rf ${SEQ_DIR}/test_1.fastq ${SEQ_DIR}/test_2.fastq --rf-stranded
kallisto quant -i ${KALLISTO_INDEX} -o ${OUT_DIR}/test_fr ${SEQ_DIR}/test_1.fastq ${SEQ_DIR}/test_2.fastq --fr-stranded

# infering strandness
paste ${OUT_DIR}/test_fr/abundance.tsv ${OUT_DIR}/test_rf/abundance.tsv ${OUT_DIR}/test_un/abundance.tsv | cut -f1,4,9,14  | awk 'BEGIN{sum1=0;sum2=0;sun3=0}{sum1+=$2;sum2+=$3;sum3+=$4}END{print sum1,sum2,sum3}' > ${OUT_DIR}/test.libtype.txt
cat ${OUT_DIR}/test.libtype.txt | awk '{print $2/$1,$3/$1,$3/$2}' | awk '{if($1<0.3 && $3>3)print "stranded";else if($1>3 && $2>3)print "reverse";else print "unstranded"}' >> ${OUT_DIR}/test.libtype.txt
```

#### Quantification of transcripts

Experiment id - GSE120561

```
kallisto quant --single --fragment-length=49 --sd=1 -b 1000 -i kallisto/index/index_trans -o kallisto/quant/w1 rawFastq/SRR7908186_W1_Worker_Pool_1.fastq.gz
kallisto quant --single --fragment-length=49 --sd=1 -b 1000 -i kallisto/index/index_trans -o kallisto/quant/w2 rawFastq/SRR7908187_W2_Worker_Pool_2.fastq.gz
kallisto quant --single --fragment-length=49 --sd=1 -b 1000 -i kallisto/index/index_trans -o kallisto/quant/aw1 rawFastq/SRR7908188_AW1_Active_Pool_1.fastq.gz
kallisto quant --single --fragment-length=49 --sd=1 -b 1000 -i kallisto/index/index_trans -o kallisto/quant/aw2 rawFastq/SRR7908189_AW2_Active_Pool_2.fastq.gz
kallisto quant --single --fragment-length=49 --sd=1 -b 1000 -i kallisto/index/index_trans -o kallisto/quant/q1 rawFastq/SRR7908190_Q1_Queen_Pool_1.fastq.gz
kallisto quant --single --fragment-length=49 --sd=1 -b 1000 -i kallisto/index/index_trans -o kallisto/quant/q2 rawFastq/SRR7908191_Q2_Queen_Pool_2.fastq.gz

#or
for i in $(ls rawFastq/); do time kallisto quant -t 4 -b 1000 -i kallisto/index/index_trans ${i} -o kallisto/quant/${i}; done
```

Experiment id - GSE93028\_GSE119256

```
SAMPLE=GSE93028_GSE119256
KALLISTO_INDEX=/mnt/c/Users/luiz_/Downloads/transcriptomeAnalysis/index/index_trans
OUT_DIR=/mnt/c/Users/luiz_/Downloads/transcriptomeAnalysis/${SAMPLE}/kallisto
SEQ_DIR=/mnt/c/Users/luiz_/Downloads/transcriptomeAnalysis/${SAMPLE}/rawFastq

time kallisto quant -t 4 -b 1000 --rf-stranded -i ${KALLISTO_INDEX} -o ${OUT_DIR}/SRR5136448 ${SEQ_DIR}/SRR5136448_V01_mRNA_RNA-Seq_1.fastq.gz ${SEQ_DIR}/SRR5136448_V01_mRNA_RNA-Seq_2.fastq.gz
time kallisto quant -t 4 -b 1000 --rf-stranded -i ${KALLISTO_INDEX} -o ${OUT_DIR}/SRR5136449 ${SEQ_DIR}/SRR5136449_V02_mRNA_RNA-Seq_1.fastq.gz ${SEQ_DIR}/SRR5136449_V02_mRNA_RNA-Seq_2.fastq.gz
time kallisto quant -t 4 -b 1000 --rf-stranded -i ${KALLISTO_INDEX} -o ${OUT_DIR}/SRR5136450 ${SEQ_DIR}/SRR5136450_V03_mRNA_RNA-Seq_1.fastq.gz ${SEQ_DIR}/SRR5136450_V03_mRNA_RNA-Seq_2.fastq.gz

time kallisto quant -t 4 -b 1000 --rf-stranded -i ${KALLISTO_INDEX} -o ${OUT_DIR}/SRR5136451 ${SEQ_DIR}/SRR5136451_Q01_mRNA_RNA-Seq_1.fastq.gz ${SEQ_DIR}/SRR5136451_Q01_mRNA_RNA-Seq_2.fastq.gz
time kallisto quant -t 4 -b 1000 --rf-stranded -i ${KALLISTO_INDEX} -o ${OUT_DIR}/SRR5136452 ${SEQ_DIR}/SRR5136452_Q02_mRNA_RNA-Seq_1.fastq.gz ${SEQ_DIR}/SRR5136452_Q02_mRNA_RNA-Seq_2.fastq.gz
time kallisto quant -t 4 -b 1000 --rf-stranded -i ${KALLISTO_INDEX} -o ${OUT_DIR}/SRR5136453 ${SEQ_DIR}/SRR5136453_Q03_mRNA_RNA-Seq_1.fastq.gz ${SEQ_DIR}/SRR5136453_Q03_mRNA_RNA-Seq_2.fastq.gz

time kallisto quant -t 4 -b 1000 --rf-stranded -i ${KALLISTO_INDEX} -o ${OUT_DIR}/SRR5136454 ${SEQ_DIR}/SRR5136454_C01_mRNA_RNA-Seq_1.fastq.gz ${SEQ_DIR}/SRR5136454_C01_mRNA_RNA-Seq_2.fastq.gz
time kallisto quant -t 4 -b 1000 --rf-stranded -i ${KALLISTO_INDEX} -o ${OUT_DIR}/SRR5136455 ${SEQ_DIR}/SRR5136455_C02_mRNA_RNA-Seq_1.fastq.gz ${SEQ_DIR}/SRR5136455_C02_mRNA_RNA-Seq_2.fastq.gz
time kallisto quant -t 4 -b 1000 --rf-stranded -i ${KALLISTO_INDEX} -o ${OUT_DIR}/SRR5136456 ${SEQ_DIR}/SRR5136456_C03_mRNA_RNA-Seq_1.fastq.gz ${SEQ_DIR}/SRR5136456_C03_mRNA_RNA-Seq_2.fastq.gz

time kallisto quant -t 4 -b 1000 --rf-stranded -i ${KALLISTO_INDEX} -o ${OUT_DIR}/SRR5136457 ${SEQ_DIR}/SRR5136457_R01_mRNA_RNA-Seq_1.fastq.gz ${SEQ_DIR}/SRR5136457_R01_mRNA_RNA-Seq_2.fastq.gz
time kallisto quant -t 4 -b 1000 --rf-stranded -i ${KALLISTO_INDEX} -o ${OUT_DIR}/SRR5136458 ${SEQ_DIR}/SRR5136458_R02_mRNA_RNA-Seq_1.fastq.gz ${SEQ_DIR}/SRR5136458_R02_mRNA_RNA-Seq_2.fastq.gz
time kallisto quant -t 4 -b 1000 --rf-stranded -i ${KALLISTO_INDEX} -o ${OUT_DIR}/SRR5136459 ${SEQ_DIR}/SRR5136459_R03_mRNA_RNA-Seq_1.fastq.gz ${SEQ_DIR}/SRR5136459_R03_mRNA_RNA-Seq_2.fastq.gz

time kallisto quant -t 4 -b 1000 --rf-stranded -i ${KALLISTO_INDEX} -o ${OUT_DIR}/SRR7770429 ${SEQ_DIR}/SRR7770429_W01_mRNA_including_lncRNA_RNA-Seq_1.fastq.gz ${SEQ_DIR}/SRR7770429_W01_mRNA_including_lncRNA_RNA-Seq_2.fastq.gz
time kallisto quant -t 4 -b 1000 --rf-stranded -i ${KALLISTO_INDEX} -o ${OUT_DIR}/SRR7770432 ${SEQ_DIR}/SRR7770432_W02_mRNA_including_lncRNA_RNA-Seq_1.fastq.gz ${SEQ_DIR}/SRR7770432_W02_mRNA_including_lncRNA_RNA-Seq_2.fastq.gz
time kallisto quant -t 4 -b 1000 --rf-stranded -i ${KALLISTO_INDEX} -o ${OUT_DIR}/SRR7770435 ${SEQ_DIR}/SRR7770435_W03_mRNA_including_lncRNA_RNA-Seq_1.fastq.gz ${SEQ_DIR}/SRR7770435_W03_mRNA_including_lncRNA_RNA-Seq_2.fastq.gz
```

Output files:

a) abundance.h5: information of bootstraps, which will be used by Sleuth.   
b) abundance.tsv: quantifications of all trancripts in the file all\_transcripts.fasta   
c) run\_info.json: informations of pseudo alignments

#### Verifying abundance values for lncov1

```
grep lncov1 kallisto/*/abundance.tsv | head
```

target\_id length eff\_length est\_counts tpm  
aw1.tsv:lncov1 1152 1104 3 1.12067  
aw2.tsv:lncov1 1152 1104 7 2.80562  
q1.tsv:lncov1 1152 1104 7 2.76799  
q2.tsv:lncov1 1152 1104 4 1.59035  
w1.tsv:lncov1 1152 1104 183 57.825  
w2.tsv:lncov1 1152 1104 153 45.2154

#### References

1 - Bray, N., Pimentel, H., Melsted, P. et al. Near-optimal probabilistic RNA-seq quantification. Nat Biotechnol 34, 525–527 (2016). https://doi.org/10.1038/nbt.3519  
2 - https://bioinfo.iric.ca/understanding-how-kallisto-works/
