## Supplemental Material 2 for "A long non-coding RNA is a key factor in the evolution of insect eusociality"

### Diferencial Expression with Sleuth

Luiz Carlos Vieira

23/03/2022

#### Sleuth for estimation of differential expression of transcripts

Sleuth is a program for differential analysis of RNA-Seq data. It makes use of quantification uncertainty estimates obtained via kallisto for accurate differential analysis of isoforms or genes, allows testing in the context of experiments with complex designs, and supports interactive exploratory data analysis via sleuth live. The sleuth methods are described in:

H Pimentel, NL Bray, S Puente, P Melsted and Lior Pachter, Differential analysis of RNA-seq incorporating quantification uncertainty, Nature Methods (2017), advanced access.

##### Instalation of leuth

```
#BiocManager::install("pachterlab/sleuth")
```

##### Loading libraries

##### Get help

```
#vignette('intro', package = 'sleuth')
```

##### Creating a list of the paths to our transcript abundance files:

First, we create a simple vector containing the paths to the directories containing the transcript abundance estimates for each sample (folders containing the .quant files).

```
base_dir <- "C:/Users/luiz_/Downloads/transcriptomeAnalysis/GSE120561/kallisto"

sample_id <- dir(file.path(base_dir))

# file.path() function gives the paths to each of the directories.
paths <- file.path(base_dir, sample_id)

paths
```

```
## [1] "C:/Users/luiz_/Downloads/transcriptomeAnalysis/GSE120561/kallisto/SRR7908186"
## [2] "C:/Users/luiz_/Downloads/transcriptomeAnalysis/GSE120561/kallisto/SRR7908187"
## [3] "C:/Users/luiz_/Downloads/transcriptomeAnalysis/GSE120561/kallisto/SRR7908188"
## [4] "C:/Users/luiz_/Downloads/transcriptomeAnalysis/GSE120561/kallisto/SRR7908189"
## [5] "C:/Users/luiz_/Downloads/transcriptomeAnalysis/GSE120561/kallisto/SRR7908190"
## [6] "C:/Users/luiz_/Downloads/transcriptomeAnalysis/GSE120561/kallisto/SRR7908191"
```

#### Creating a metadata dataframe

Create metadata associated with the kallisto files using the data.frame().

```
# Sleuth requires a column entitled "sample" containing the sample names:
metadata <- data.frame(sample = sample_id,
                       condition = factor(c(rep("W", 2),
                                            rep("AW", 2),
                                            rep("Q", 2))))
```

#### Naming the vector of directory paths with the corresponding sample names

```
# Name the directory paths for the abundance files with their corresponding sample IDs
names(paths) <- sample_id

paths
```

```
##                                                                      SRR7908186
## "C:/Users/luiz_/Downloads/transcriptomeAnalysis/GSE120561/kallisto/SRR7908186"
##                                                                      SRR7908187
## "C:/Users/luiz_/Downloads/transcriptomeAnalysis/GSE120561/kallisto/SRR7908187"
##                                                                      SRR7908188
## "C:/Users/luiz_/Downloads/transcriptomeAnalysis/GSE120561/kallisto/SRR7908188"
##                                                                      SRR7908189
## "C:/Users/luiz_/Downloads/transcriptomeAnalysis/GSE120561/kallisto/SRR7908189"
##                                                                      SRR7908190
## "C:/Users/luiz_/Downloads/transcriptomeAnalysis/GSE120561/kallisto/SRR7908190"
##                                                                      SRR7908191
## "C:/Users/luiz_/Downloads/transcriptomeAnalysis/GSE120561/kallisto/SRR7908191"
```

#### Combining the metadata with the paths

Combining the metadata with the paths to the transcript abundance files to use as input for the Sleuth analysis.

Sleuth requires a column entitled "path" containing the paths to the estimated counts files stored in our sf_dirs:

```
# Adding a column named 'path'
metadata$path <- paths

metadata
```

```
##       sample condition
## 1 SRR7908186         W
## 2 SRR7908187         W
## 3 SRR7908188        AW
## 4 SRR7908189        AW
## 5 SRR7908190         Q
## 6 SRR7908191         Q
##                                                                     path
## 1 C:/Users/luiz_/Downloads/transcriptomeAnalysis/GSE120561/kallisto/SRR7908186
## 2 C:/Users/luiz_/Downloads/transcriptomeAnalysis/GSE120561/kallisto/SRR7908187
## 3 C:/Users/luiz_/Downloads/transcriptomeAnalysis/GSE120561/kallisto/SRR7908188
## 4 C:/Users/luiz_/Downloads/transcriptomeAnalysis/GSE120561/kallisto/SRR7908189
## 5 C:/Users/luiz_/Downloads/transcriptomeAnalysis/GSE120561/kallisto/SRR7908190
```

```
## 6 C:/Users/luiz_/Downloads/transcriptomeAnalysis/GSE120561/kallisto/SRR7908191
```

##### Defining condition levels

```r
metadata$condition <- relevel(metadata$condition, ref = "W")
```

##### Creating a variable containing the model design

```r
design <- ~ condition
```

#### Create sleuth object for analysis

```r
so <- sleuth_prep(metadata,
                  full_model = design,
                  #target_mapping = t2g,
                  num_cores = 4L,
                  read_bootstrap_tpm = TRUE,
                  extra_bootstrap_summary = TRUE,
                  transform_fun_counts = function(x) log2(x + 0.5)
                  )
```

```
## Warning in check_num_cores(num_cores): It appears that you are running Sleuth from within Rstudio.
## Because of concerns with forking processes from a GUI, 'num_cores' is being set to 1.
## If you wish to take advantage of multiple cores, please consider running sleuth from the command lin

## reading in kallisto results

## dropping unused factor levels

## ......
## normalizing est_counts
## 17599 targets passed the filter
## normalizing tpm
## merging in metadata
## summarizing bootstraps
## ......
```

NOTE: By default the transformation of counts is natural log, which would make the output fold changes somewhat more difficult to interpret. By specifying the transform_fun_counts to be log2(x + 0.5) we are ensuring our output fold changes are log2.

##### Fitting the sleuth model

```r
so <- sleuth_fit(so)
```

```
## fitting measurement error models

## shrinkage estimation

## computing variance of betas
```

##### Check which models have been fit and which coefficients can be tested

```r
models(so)
```

```
## [  full  ]
## formula:  ~condition
## data modeled:  obs_counts
## transform sync'ed:  TRUE
## coefficients:
##  (Intercept)
##      conditionAW
##      conditionQ
```

##### Test significant differences between conditions using the Wald test

```
DE_AW <- sleuth_wt(so, which_beta = 'conditionAW')


DE_Q <- sleuth_wt(so, which_beta = 'conditionQ')


# Get results
sleuth_results_AW <- sleuth_results(DE_AW,
                                    test = 'conditionAW',
                                    show_all = TRUE)

sleuth_results_Q <- sleuth_results(DE_Q,
                                   test = 'conditionQ',
                                   show_all = TRUE)
```

##### Salving results table

```
#write.xlsx(sleuth_results_AW, file='results/sleuth_results_AW.xlsx', sheetName = "DE_AW",
#  col.names = TRUE, row.names = FALSE, append = FALSE)
#
#write.xlsx(sleuth_results_Q, file='results/sleuth_results_Q.xlsx', sheetName = "DE_Q",
#  col.names = TRUE, row.names = FALSE, append = FALSE)
```

##### Visualization of resultus with shiny app.

```
#sleuth_live(so)
```

---

---

#### Results and visualization

---

---

List of transcrits of interest: "XM_001120691.5", "XM_393605.7", "XM_624635.6", "lncov1"

```
txI <- c("XM_001120691.5", "XM_393605.7", "XM_624635.6", "lncov1")
```

#### Get Differencial expressed transcripts results table

```
sleuth_sig_AW <- dplyr::filter(sleuth_results_AW, pval <= 0.05)
sleuth_sig_Q <- dplyr::filter(sleuth_results_Q, pval <= 0.05)

head(sleuth_sig_AW)
```

```
##        target_id          pval          qval         b      se_b  mean_obs
## 1 XM_001120991.5  0.000000e+00  0.000000e+00 -5.666737 0.1399872 10.047758
## 2 XM_026440745.1 2.611400e-211 2.297902e-207 -4.523997 0.1458256  9.859082
## 3    XM_392190.7 4.276566e-191 2.508776e-187 -6.459788 0.2190754  9.805255
## 4 XM_003250895.4 3.014828e-184 1.326449e-180  7.613183 0.2629994  8.950848
## 5 XM_001122993.5 2.111095e-163 7.430632e-160 -4.420064 0.1622564  8.603663
## 6 XM_016918022.2 6.760192e-159 1.982877e-155  3.705620 0.1379696 10.485286
##     var_obs    tech_var     sigma_sq smooth_sigma_sq final_sigma_sq
## 1 10.922706 0.007051827  0.008617699      0.01254458     0.01254458
## 2  8.252392 0.007326495  0.013938605      0.01231857     0.01393860
## 3 13.648660 0.010117252  0.037876769      0.01228386     0.03787677
## 4 15.724869 0.044848930  0.024319764      0.01337651     0.02431976
## 5  5.127883 0.011293629 -0.001990372      0.01503351     0.01503351
## 6  4.730991 0.003443490  0.015592129      0.01374573     0.01559213
```

#### Get bootstrap summary

Getting the Maximum likelihood estimation est_counts or tpm from bootstraps from all samples.

```
head(as.data.frame(so$obs_norm), 10)
```

```
##         target_id     sample  est_counts         tpm eff_len  len
## 1          lncov1 SRR7908186  160.086820   58.786597    1104 1152
## 2          lncov1 SRR7908187  134.803328   49.070471    1104 1152
## 3          lncov1 SRR7908188    2.986359    1.072901    1104 1152
## 4          lncov1 SRR7908189    7.486275    2.688563    1104 1152
## 5          lncov1 SRR7908190    7.691013    2.746850    1104 1152
## 6          lncov1 SRR7908191    4.436825    1.583259    1104 1152
## 7  NM_001010975.1 SRR7908186 5108.781579 1721.644664    1203 1251
## 8  NM_001010975.1 SRR7908187 7028.275494 2347.858054    1203 1251
## 9  NM_001010975.1 SRR7908188 3818.557944 1258.984145    1203 1251
## 10 NM_001010975.1 SRR7908189 3284.335924 1082.443865    1203 1251
```

Getting the bootstraps summary of est_counts or tpm from each sample.

```
s1 <- as.data.frame(so$bs_quants$SRR7908186$est_counts)

dplyr::filter(s1, row.names(s1) == "lncov1")
```

```
##             min    lower     mid    upper      max
## lncov1 6.982651 7.254686 7.32721 7.388751 7.662325
```

Getting the bootstrap summary from all samples with the function get_bootstrap_summary()

```
bt_summ <- get_bootstrap_summary(so, "lncov1", units = "est_counts")
bt_summ
```

```
##                  min    lower      mid    upper      max     sample condition
## SRR7908186  6.982651 7.254686 7.327210 7.388751 7.662325 SRR7908186         W
## SRR7908187  6.694670 7.002869 7.089418 7.162219 7.428847 SRR7908187         W
## SRR7908188 -1.000000 1.316671 1.801721 2.164082 3.517274 SRR7908188        AW
```

```
## SRR7908189 -1.000000 2.547780 2.997523 3.339880 4.223496 SRR7908189         AW
## SRR7908190 -1.000000 2.583418 3.034042 3.376908 4.176279 SRR7908190          Q
## SRR7908191 -1.000000 1.936447 2.303583 2.595988 3.899145 SRR7908191          Q
```

#### Volcano plot

volcano plot, Plots of beta value (regression) versus log of significance p-values.

##### Volcano plot Queen vs Worker

```
q <- as.data.frame(sleuth_results_Q)
q <- na.omit(q)
q$significant <- ifelse(q$pval<0.05, "True", "False")
q[which(abs(q$b)<0.5),'significant'] <- "False"
q <- q[order(q$pval),]


legen <- q[q$target_id %in% txI, ]


volcano = ggplot(q, aes(b, -log10(pval))) +
  geom_point(aes(col=significant)) +
  scale_color_manual(values=c("red", "gray"))


volcano + geom_label_repel(data= legen, aes(label= legen$target_id), size=4,
  box.padding = unit(2, "lines"), point.padding = unit(5, "points")) +
  labs(title= "Volcano plot of Diferencial Expressed Transcripts",
       subtitle = "Comparison of Queen vs Worker",
       x= "log2 fold change",
       y= "p-value (-log10)",
       color="Differentially Expressed") +
  theme_bw() + coord_cartesian(clip = "off")
```

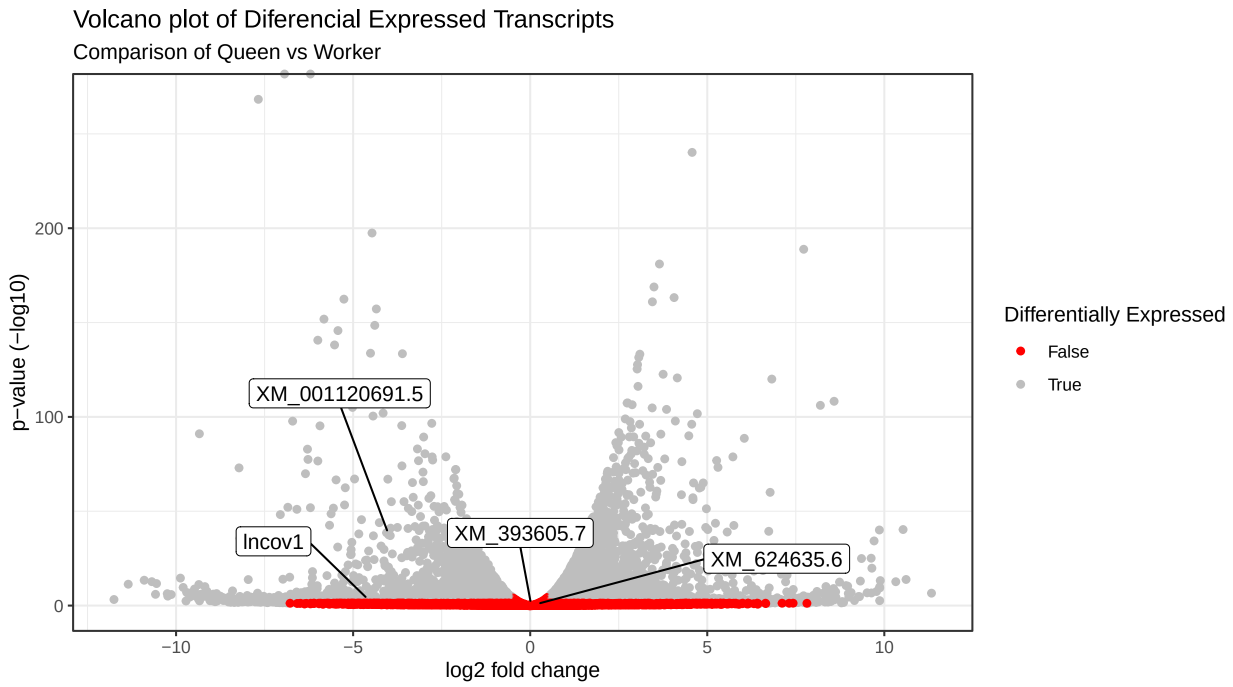


##### Volcano plot Queenless-worker vs Worker

```
q <- as.data.frame(sleuth_results_AW)
q <- na.omit(q)
q$significant <- ifelse(q$pval<0.05, "True", "False")
q[which(abs(q$b)<0.5),'significant'] <- "False"
q <- q[order(q$pval),]


legen <- q[q$target_id %in% txI, ]


volcano = ggplot(q, aes(b, -log10(pval))) +
  geom_point(aes(col=significant)) +
  scale_color_manual(values=c("red", "gray"))


volcano + geom_label_repel(data= legen, aes(label= legen$target_id), size=4,
  box.padding = unit(2, "lines"), point.padding = unit(5, "points")) +
  labs(title= "Volcano plot of Diferencial Expressed Transcipts",
       subtitle = "Comparison of Queenless-Worker vs Worker",
       x= "log2 fold change",
       y= "p-value (-log10)",
       color="Differentially Expressed") +
  theme_bw() + coord_cartesian(clip = "off")
```

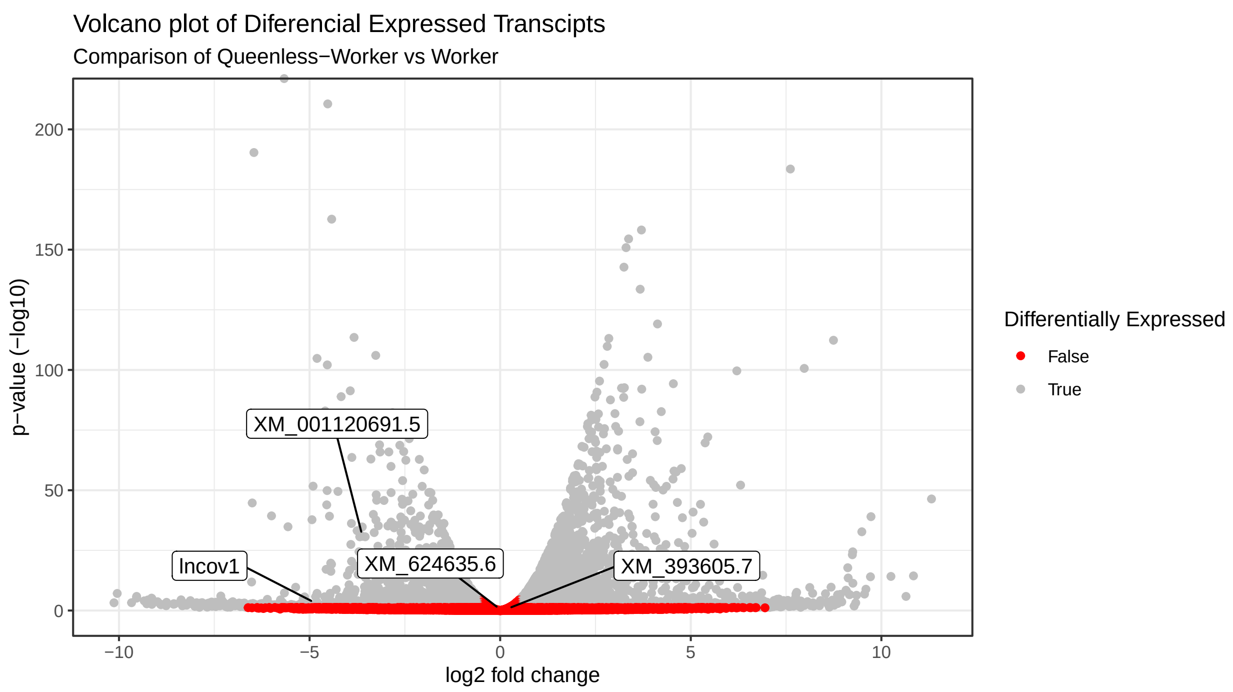


#### PCA

```r
#png("pca-transcritos.png",height = 8, width = 10, units = 'in',res=600)

plot_pca(so, pc_x = 1L, pc_y = 2L, use_filtered = TRUE,
  units = "est_counts", text_labels = F, color_by = "condition",
  point_size = 4) +
  theme_bw() + # remove default ggplot2 theme
  ggtitle(label = "Principal Component Analysis (PCA)",
          subtitle = "Variance among worker, queenless-worker and queen")
```

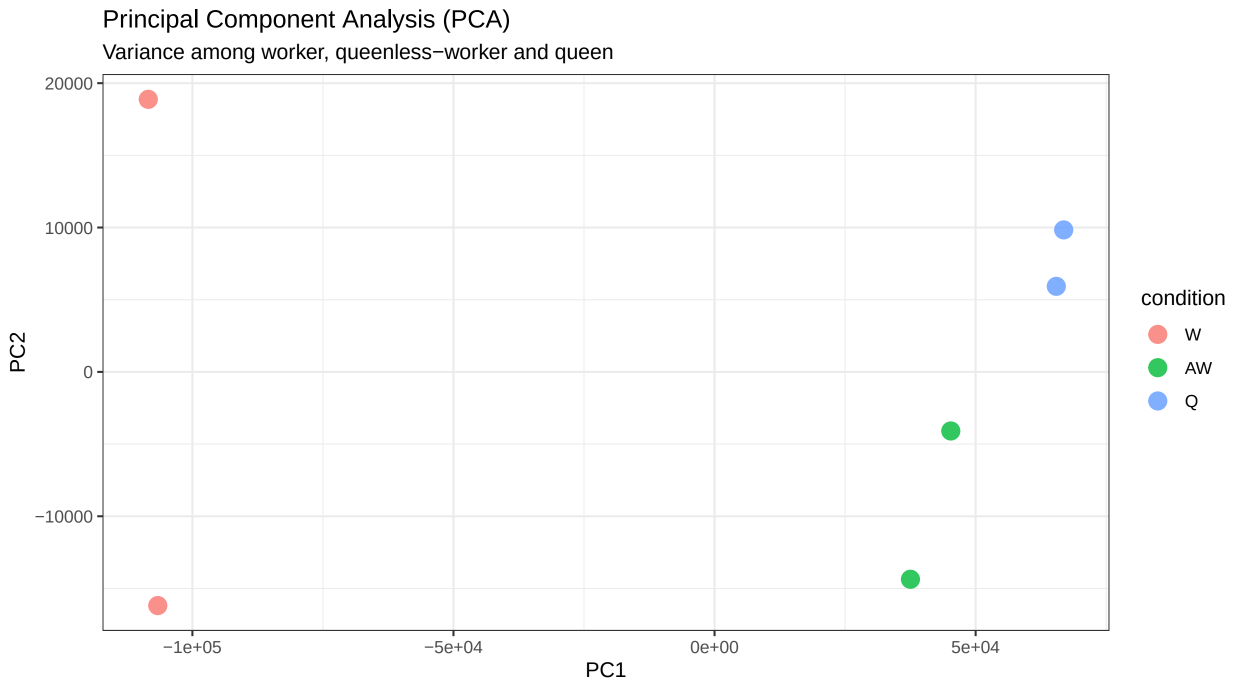


#### Density

```r
plot_group_density(so, use_filtered = TRUE, units = "est_counts",
                   trans = "log", grouping = "condition", offset = 1)
```

```
## Warning: `group_by_()` was deprecated in dplyr 0.7.0.
## Please use `group_by()` instead.
## See vignette('programming') for more help
## This warning is displayed once every 8 hours.
## Call `lifecycle::last_lifecycle_warnings()` to see where this warning was generated.
```

```
## Warning: `summarise_()` was deprecated in dplyr 0.7.0.
## Please use `summarise()` instead.
## This warning is displayed once every 8 hours.
## Call `lifecycle::last_lifecycle_warnings()` to see where this warning was generated.
```

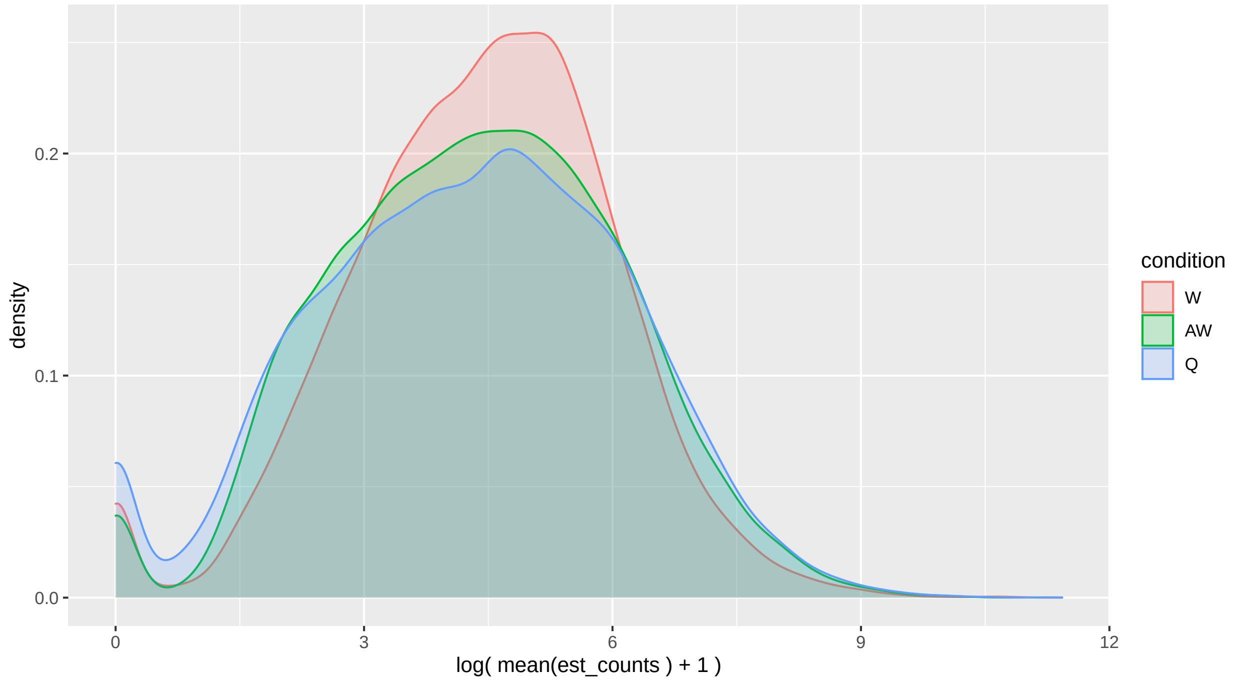


##### Boxplots

Transcript lncov1

```
names_facet <- c(W = "Queenright Workers", AW = "Queenless Workers", Q = "Queen")

plot1 <- plot_bootstrap(so, "lncov1", units = "est_counts", color_by = "condition") +

ggtitle("lncov1") + theme_bw() +
  theme(legend.position="none", plot.title = element_text(size = 14, face="italic")) +
  ylab(expression("normalized read counts - log"[2])) + xlab("Samples") +
  facet_rep_wrap(~condition, labeller = as_labeller(names_facet), scales='free_x', repeat.tick.labels =

plot1
```

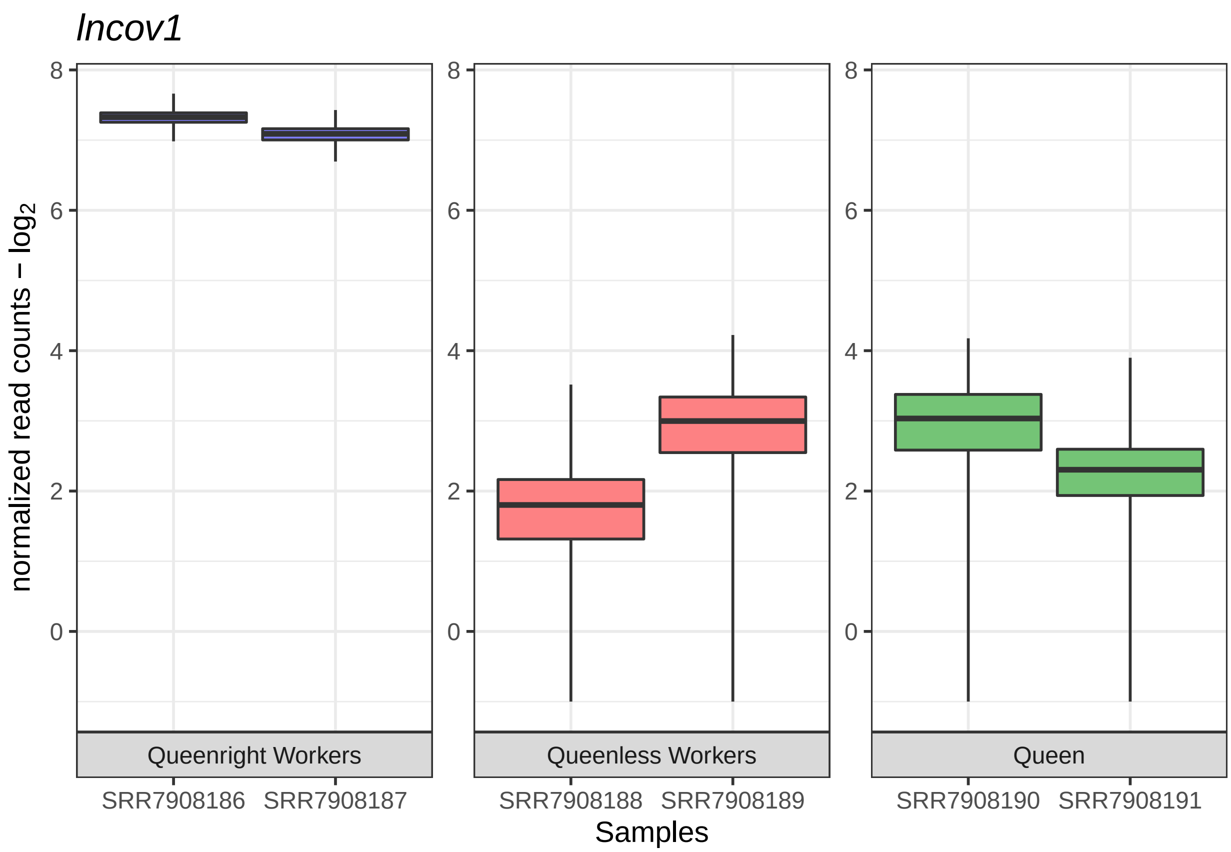


Transcript XM_001120691.5 = Loc726407

```
plot2 <- plot_bootstrap(so, "XM_001120691.5", units = "est_counts", color_by = "condition") +

ggtitle("Loc726407") + theme_bw() + theme(legend.position="none", plot.title = element_text(size = 14, f
  ylab(expression("normalized read counts - log"[2])) + xlab("Samples") +
  facet_rep_wrap(~condition, labeller = as_labeller(names_facet), scales='free_x', repeat.tick.labels =
  scale_fill_manual(values=c("#7f7efd", "#fd8183", "#74C476"))

plot2
```

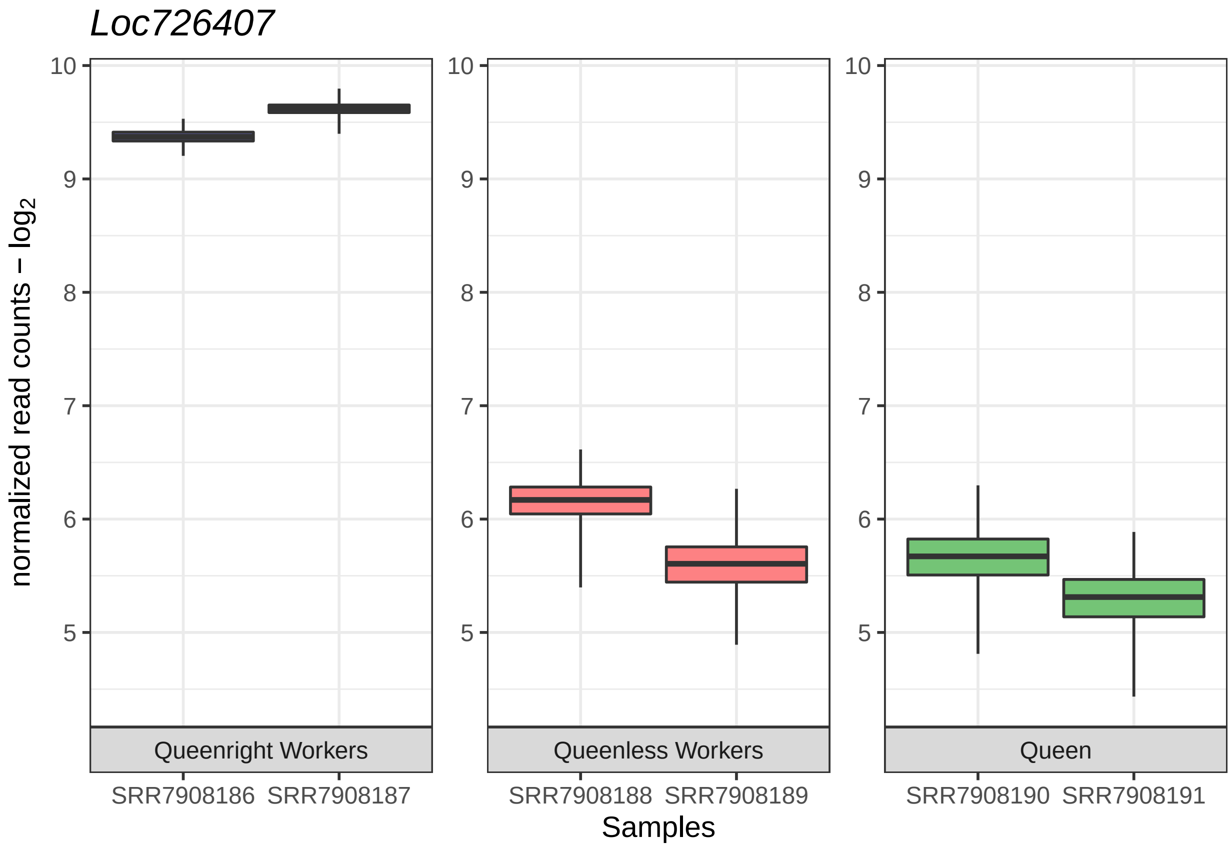


Transcript XM_393605.7 = Gapdh

```
plot3 <- plot_bootstrap(so, "XM_393605.7", units = "est_counts", color_by = "condition") +

ggtitle("Gapdh") + theme_bw() + theme(legend.position="none", plot.title = element_text(size = 14, face=
  ylab(expression("normalized read counts - log"[2])) + xlab("Samples") +
  facet_rep_wrap(~condition, labeller = as_labeller(names_facet), scales='free_x', repeat.tick.labels =
  scale_fill_manual(values=c("#7f7efd", "#fd8183", "#74C476"))

plot3
```

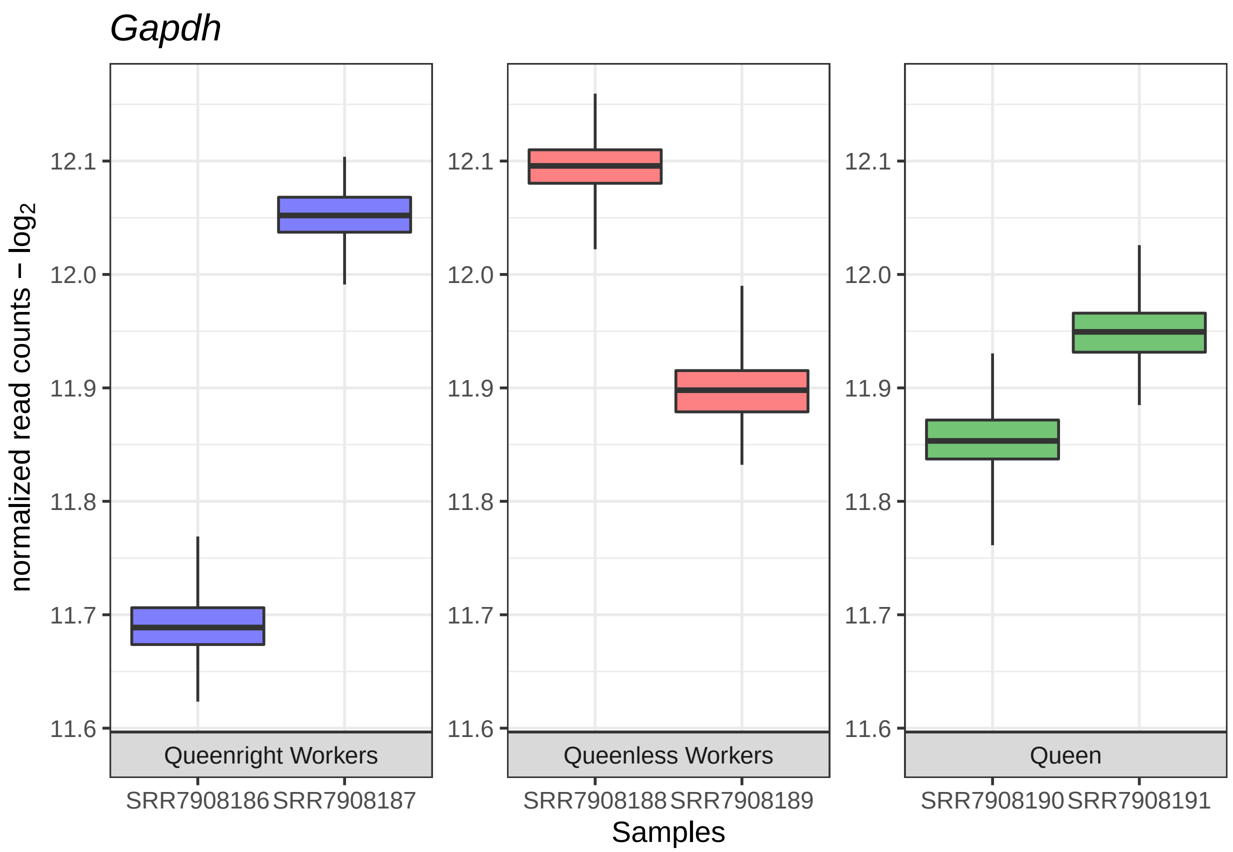


XM_624635.6 para Tudor-SN

```
plot4 <- plot_bootstrap(so, "XM_624635.6", units = "est_counts", color_by = "condition") +

ggtitle("Tudor-SN") + theme_bw() +
  theme(legend.position="none", plot.title = element_text(size = 14, face="italic")) +
  ylab(expression("normalized read counts - log"[2])) + ylim(c(10, 10.75)) + xlab("Samples") +
  facet_rep_wrap(~condition, labeller = as_labeller(names_facet), scales='free_x', repeat.tick.labels =
  scale_fill_manual(values=c("#7f7efd", "#fd8183", "#74C476"))

plot4
```

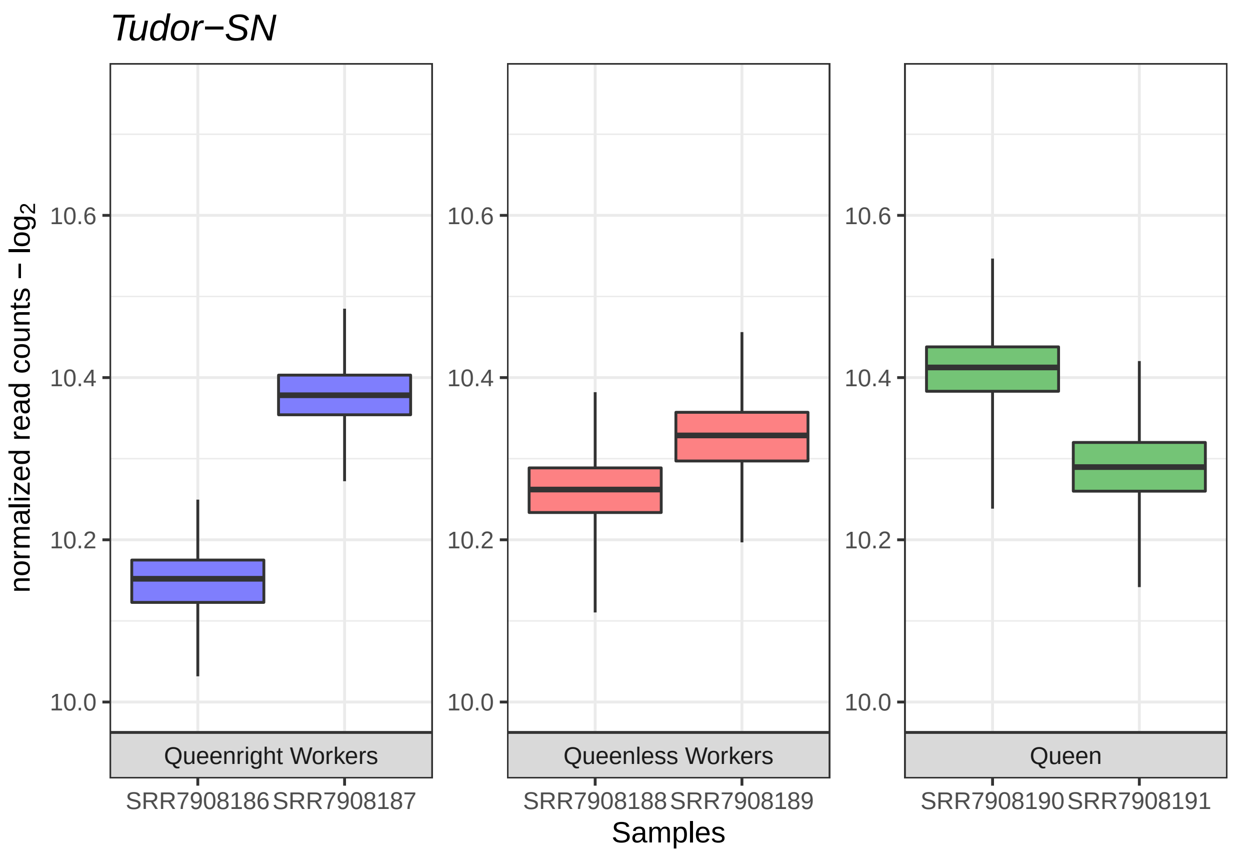


Distribuition of est_counts of transcripts:

```
plot_grid(plot1 + theme(legend.position="right"),
          plot2 + theme(legend.position="right"),
          plot3 + theme(legend.position="right"),
          plot4 + theme(legend.position="right"),
          #plot4 + theme(legend.position="none") + ylab(NULL),
          labels = c("A", "B", "C", "D"),
          ncol = 1, nrow = 4)
```

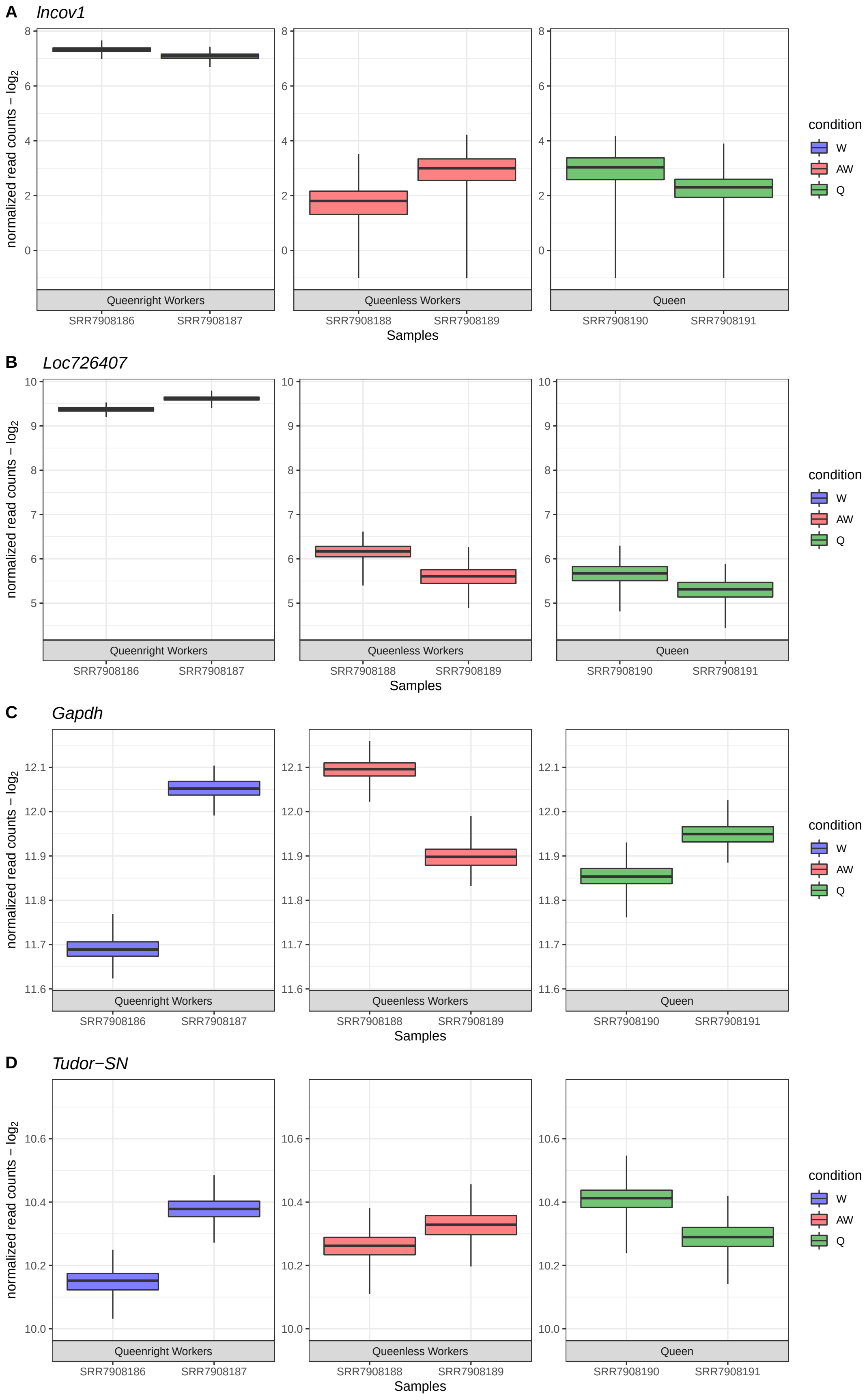
A
lncov1
B
Loc726407
C
Gapdh
D
Tudor−SN
normalized read counts − $log_2$
Queenright Workers
Queenless Workers
Queen
SRR7908186
SRR7908187
SRR7908188
SRR7908189
SRR7908190
SRR7908191
Samples
condition
W
AW
Q

#### 6. Heatmaps

we can perform an expression heatmap for select transcripts, like transcripts of interest:

```
mat <- sleuth_results_Q[sleuth_results_Q$target_id %in% txI, ]

plot_transcript_heatmap(DE_Q, units = "est_counts", transcripts = mat$target_id)
```

#### Referências


https://pachterlab.github.io/sleuth/walkthroughs
https://pachterlab.github.io/sleuth_walkthroughs/trapnell/analysis.html
https://www.nature.com/articles/nmeth.4324.pdf?origin=ppub
https://liorpachter.wordpress.com/2015/08/17/a-sleuth-for-rna-seq/
https://hbctraining.github.io/DGE_workshop_salmon/lessons/09_sleuth.html#:~:text=What%20is%20Sleuth%3F,expression
https://rdrr.io/github/pachterlab/sleuth/src/R/plots.R
https://www.simplypsychology.org/boxplots.html
